# Origin of electroneutrality in living system

**DOI:** 10.1101/2021.09.21.461264

**Authors:** Amir Akbari, Bernhard O. Palsson

**Author notes:** Email addresses:* (Amir Akbari), (Bernhard O. Palsson).

## Abstract

Identifying the first chemical transformations, from which life emerged is a central problem in the theories of life’s origins. These reactions would likely have been self-sustaining and self-reproductive before the advent of complex biochemical pathways found in modern organisms to synthesize lipid membranes, enzymes, or nucleic acids. Without lipid membranes and enzymes, exceedingly low concentrations of the organic intermediates of early metabolic cycles in protocells would have significantly hindered evolvability. To address this problem, we propose a mechanism, where a positive membrane potential elevates the concentration of the organic intermediates. In this mechanism, positively charged surfaces of protocell membranes due to accumulation of transition metals generate positive membrane potentials. We compute steady-state ion distributions and determine their stability in a protocell model to identify the key factors constraining achievable membrane potentials. We find that (i) violation of electroneutrality is necessary to induce nonzero membrane potentials; (ii) strategies that generate larger membrane potentials can destabilize ion distributions; and (iii) violation of electroneutrality enhances osmotic pressure and diminishes reaction efficiency, thereby driving the evolution of lipid membranes, specialized ion channels, and active transport systems.

**Significance:** The building blocks of life are constantly synthesized and broken down through concurrent cycles of chemical transformations. Tracing these reactions back 4 billion years to their origins has been a long-standing goal of evolutionary biology. The first metabolic cycles at the origin of life must have overcome several obstacles to spontaneously start and sustain their nonequilibrium states. Notably, maintaining the concentration of organic intermediates at high levels needed to support their continued operation and subsequent evolution would have been particularly challenging in primitive cells lacking evolutionarily tuned lipid membranes and enzymes. Here, we propose a mechanism, in which the concentration of organic intermediates could have been elevated to drive early metabolic cycles forward in primitive cells with ion-permeable porous membranes under prebiotic conditions and demonstrate its feasibility in a protocell model from first principles.

## Introduction

How complex life originated on a lifeless planet from nothing but a few inorganic precursors is a mystery [1]. Darwin postulated that evolution is a slow, stepwise process involving a gradual accumulation of small variations over geological time scales [2]. From this viewpoint, the structure of extant biology could still hold relics of the first life forms on the primitive Earth [3]. A striking feature of extant life is particularly relevant in this regard: All anabolic and catabolic processes in modern organisms proceed through five universal intermediates, all involved in the oxidative and reductive tri-carboxylic acid cycle [4]. Therefore, it is plausible to assume that all of biochemistry, as we know it, emerged from a set of most ancient chemical transformations involving the five universal intermediates that could have operated and evolved under prebiotic conditions. This is, in fact, one of the main paradigms in the origins-of-life literature, known as the metabolism-first hypothesis [5, 6, 7, 8]. It assumes that life originated from primitive metabolic reactions that could have spontaneously started and functioned without relying on molecular apparatuses, such as organic cofactors, lipid bilayers, enzymes, or genetic replication that are characteristics of modern cells.

As with other origins-of-life theories, metabolism-first theories must address a key challenge to explain abiogenesis: provide a plausible explanation for how prebiotic chemistry transitioned into biochemistry without sophisticated molecular machinery that all living systems depend on today. Notably, organic molecules could not have been contained in compartments without lipid membranes. Electroneutrality, metal-ion homeostasis, and pH homeostasis could not have been achieved without specialized ion channels. A continuous supply of energy could not have been provided to fuel metabolism without membrane proteins, and metabolic reactions could not have proceeded fast enough to synthesize necessary precursors for the evolution of complex reaction networks without enzymes or genetic replication.

In this article, we focus on one of the main obstacles that first metabolic cycles must have overcome to evolve within a metabolism-first view of life’s origins: to elevate the concentration of organic intermediates in primitive cells lacking lipid membranes and enzymes beyond a minimum level required to drive the metabolic cycles forward. The concentration of these intermediates would have been set by a balance between two counteracting mass fluxes, namely membrane transport and reaction rate. The transport rate of organic molecules through porous membranes of primitive cells would have been significantly higher than through lipid bilayers in modern cells, while the rates of nonenzymatic reactions would have been much lower than enzymatic counterparts. Therefore, maintaining the concentration of organic molecules in primitive cells at levels comparable to modern organisms would have been difficult, if not impossible.

We examine the foregoing problem more closely by developing a protocell model of life’s origins to understand how the membrane potential and electroneutrality could have influenced the emergence of the earliest metabolic cycles. Specifically, we seek to answer (i) whether a positive membrane potential could have developed across protocell membranes under steady state conditions, large enough to concentrate negatively charged organic intermediates of early metabolic networks inside primitive cells, and (ii) if the steady-state solutions could have been stable.

## Results

### Model Description

To examine possible mechanisms that could have helped concentrate the organic intermediates of first metabolic cycles inside primitive cells, we consider a spherical cell of radius *R*_*c*_ with an ion-permeable porous membrane of thickness *d*, residing in the primitive ocean (Fig. 1A). The ocean is assumed electroneutral, only comprising monovalent salts. For simplicity, we only consider two salts with concentrations 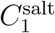 and 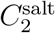. Hereafter, we refer to these salts as salt-I and salt-II. Dissociation of each salt in water yields equal amounts of its constituent cation and anion. Thus, the total concentration of the resulting ions in the ocean is *C*_*∞*_ = 2*C*^salt^ with 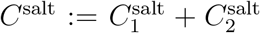 the total salt concentration (see “SI: Protocell Model” for details). Ions can transfer between the cell and ocean due to a concentration gradient or nonuniform electric-potential field induced by an uneven distribution of cations and anions in the space.

**Figure 1:**
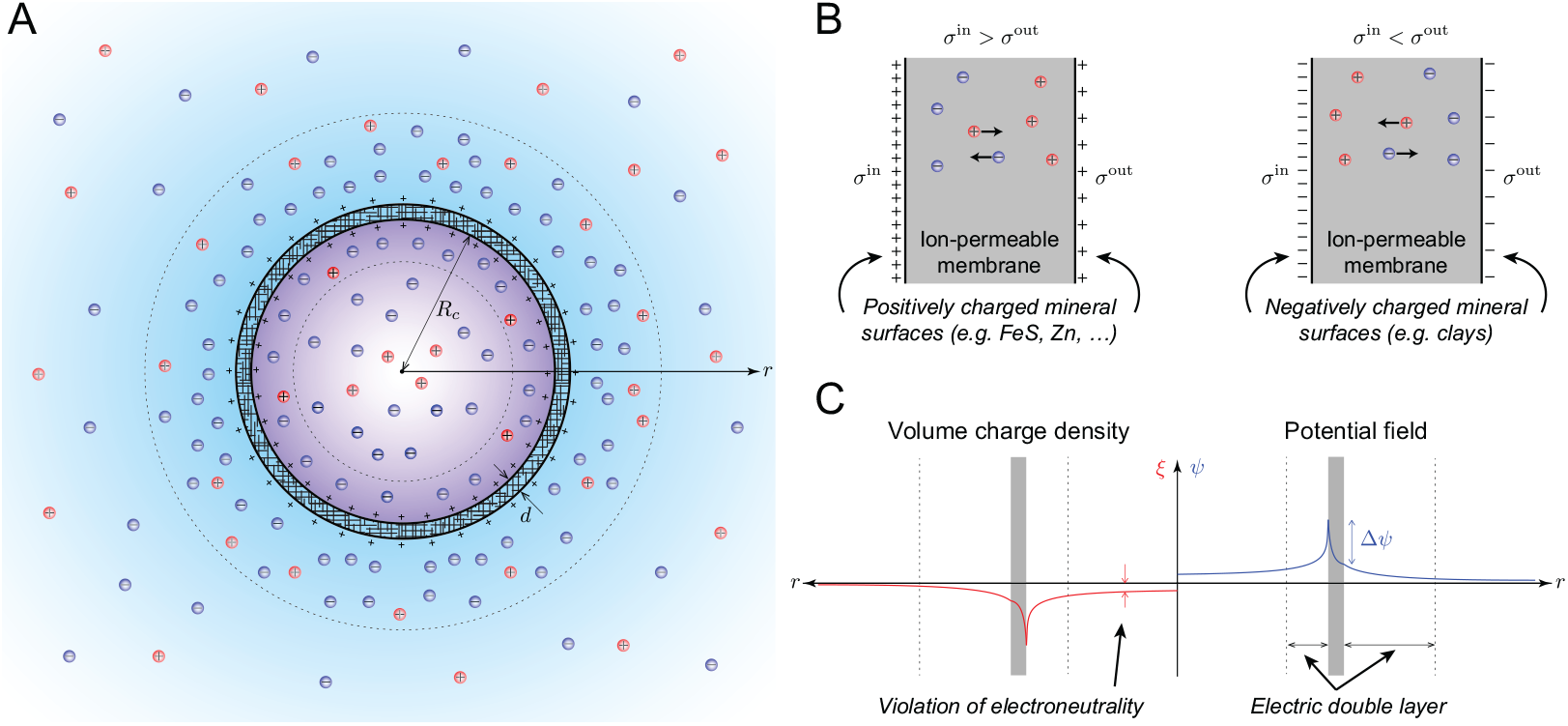
Proposed protocell model to study the origin of the membrane potential and electroneutrality. (A) Schematic representation of charge distribution and electric double layers that could have developed inside and around protocells with ion-permeable membranes at the bottom of the primitive ocean. The inner and outer surfaces of the membrane could have been positively charged due to the presence of transition-metal minerals. (B) Two possible mechanisms, through which a nonzero membrane potential could have developed across protocell membranes. The left diagram shows a case, where the inner and outer surfaces of the membrane are positively charged, resulting in a positive membrane potential. The right diagram shows the opposite case, where the inner and outer surfaces are negatively charged, resulting in a negative membrane potential. In both cases, the magnitude of the surface charge density on the inner surface *σ*^in^ is assumed to be larger than that on the outer surface *σ*^out^. (C) Possible radial profiles for the electric potential *ψ* and volume charge density *ξ* that could lead to a positive membrane potential. Here, the electroneutrality constraint inside the cell and membrane is globally relaxed to achieve a positive membrane potential.

Note that, our goal here is not to computationally exhaust all possible scenarios and physico-chemical systems that could have brought about life on the primitive Earth. Rather, we aim to study a tractable and plausible model that possesses the most essential features of primitive cells to gain qualitative insights into the restrictiveness of the constraints on primordial metabolic evolution through concrete and quantitative analyses.

The protocell model described above is a simplified version of how primitive cells might have operated in the early ocean. A key simplification concerns the composition of the primitive ocean with regards to the number and valence of ions. As evidenced by leaching studies of oceanic crusts, early oceans were likely more saline than modern seawater, comprising several monovalent and polyvalent ions in high concentrations (*e.g.*, Na^+^, K^+^, Ca^2+^, 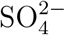) [9]. Nevertheless, by understanding the fundamental constraints that govern electric-potential and charge distributions in our simplified model, we may find clues to possible mechanisms, through which a positive membrane potential could have been achieved in the general case.

In modern cells, there are two main constraints that govern the flow of ions into and out of the cell, namely species mass balance and electroneutrality [10, 11]. Thanks to several specialized ion channels in their ion-impermeable lipid membranes, modern cells can generate local ion gradients in their membranes and a nonzero membrane potential while maintaining electroneutrality on both sides of the membrane [12]. However, in ion-permeable protocell membranes with no specialized ion channels, was it possible to generate a nonzero membrane potential without violating electroneutrality inside the cell? If not, what forms of charge distribution could have furnished a nonzero membrane potential? Was it necessary for electroneutrality to be violated locally or globally? What mechanism could have underlain electroneutrality violation?

To answer these questions, we replace electroneutrality with Maxwell’s first law—a more fundamental constraint governing the interdependence of charge and electric potential

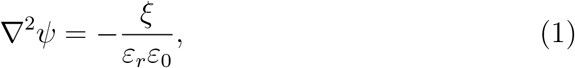

where *ψ* is the electric potential, *ξ* volume charge density, *ε*_*r*_ relative permittivity of the medium (water in our model), and *ε*_0_ vacuum permittivity. This equation describes the relationship between charge and electric-potential distribution in the cell, membrane, and ocean, whether or not electroneutrality holds. The species mass balance equation reads

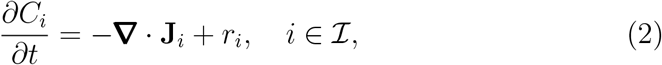

where 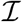 is the index set of all the species involved in the system with *C*_*i*_, **J**_*i*_, and *r*_*i*_ the concentration, flux vector, and production rate of species *i*. We determine the steady-state solutions of Eqs. (1) and (2) in the cell, membrane, and ocean to identify the key parameters affecting the electric potential distribution (see “SI: Governing Equations”). We then, through numerical experiments, determine what parameter values lead to a positive membrane potential, and, if so, whether electroneutrality can be maintained.

We note that Eq. (2) can be simplified by decoupling the mass balance equation for reactive (*i.e.*, inorganic cofactors, reducing agents, energy sources, and organic intermediates of metabolic cycles) and nonreactive (*i.e.*, inorganic ions of the primitive ocean) species. The earliest metabolic reactions are believed to have been catalyzed by naturally occurring minerals at a significantly smaller rate than their enzymatic counterparts [13, 8, 14, 15]. As a result, the concentration of reactive compounds produced by these reactions would have been extremely small [14, 16], especially compared to that of inorganic ions in the primitive ocean [9]. Therefore, reactive species could not have significantly contributed to the development of the membrane potential. Accordingly, we neglect their contribution in Eqs. (1) and (2), solving Eq. (2) only for inorganic ions. These ions are not consumed or produced by any metabolic reactions taking place inside the protocell, so that they attain their steady state much faster than reactive species. Therefore, we solve Eq. (2) for 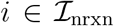, where *r*_*i*_ = 0 with 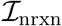 the index set of nonreactive species.

Given that the ocean is electroneutral in our model, one may deduce from Maxwell’s first law that inducing a nontrivial electric potential field in the cell and membrane is not possible without any charge sources. We, thus, consider a scenario, where charged mineral species cover the inner and outer surfaces of the membrane, giving rise to surface charge densities *σ*^in^ and *σ*^out^ (Fig. 1B). Transition-metal and clay minerals tend to have positively and negatively charged surfaces, respectively. Here, the idea is that the difference between *σ*^in^ and *σ*^out^ could generate a nontrivial electric potential field and a nonzero membrane potential as a result. For example, suppose that |*σ*^in^| > |>*σ*^out^|. Then, positively charged mineral surfaces can generate a positive membrane potential (Fig. 1C), and negatively charged mineral surfaces can generate a negative membrane potential.

Interestingly, the foregoing scenario is consistent with Wächtershäuser’s theory of surface metabolism [5]. In this theory, positively charged mineral surfaces, such as those of divalent transition metals, play a central role in facilitating early metabolic reactions because of the strong ionic bonds that form between negatively charged organic molecules and transition metals. In our model, transition metals not only catalyze early metabolic reactions, but also can help concentrate organic molecules inside primitive cells by generating a positive membrane potential if deposited on the surfaces of protocell membranes.

### Positive Membrane Surface Charges Induce Positive Membrane Potential

We first studied the steady-state solutions of Eqs. (1) and (2), assuming a spherical symmetry. From our preliminary numerical experiments, we identified three key parameters most affecting the membrane potential, namely the total salt concentration in the ocean *C*^salt^, membrane surface charge density *σ*, and cell radius *R*_*c*_. We then constructed parametric steady-state solutions with respect to these parameters. Solutions were computed numerically using fourth-order finite-difference schemes (see “SI: Numerical Approximation of Steady-State Solutions”). In this section, we present steady-state solutions parametrized with *C*^salt^ at fixed *R*_*c*_, *σ*, and *σ*_*r*_, where *σ*^in^ = *σ*, and *σ*^out^ = *σ*_*r*_*σ*. All other parameters used to obtain the results in this and subsequent sections are summarized in Table S1.

Steady-state solutions of the electric potential, volume charge density, and ion concentrations in the cell, membrane, and ocean exhibit a qualitatively similar trend for all values of *C*^salt^ (Fig. 2). Positively charged membrane surfaces induce a positive electric potential field in the entire domain (0 ≤ *r <* ∞) (Fig. 2A). The resulting membrane potential is also positive, scaling inversely with *C*^salt^. Electric double layers form around the inner and outer surfaces of the membrane due to surfaces charges. Consequently, negative ions concentrate inside the electric double layers (Figs. 2C,D), leading to a net negative volume charge density throughout the entire domain (Fig. 2B). We also performed the same analysis for smaller *σ* and found that the electric potential, volume charge density, and the membrane potential all tend to vanish as *σ* → 0. Overall, our results suggest that membrane surface charges generate a nonzero membrane potential. Electroneutrality is then violated in the cell, membrane, and ocean as a consequence.

**Figure 2:**
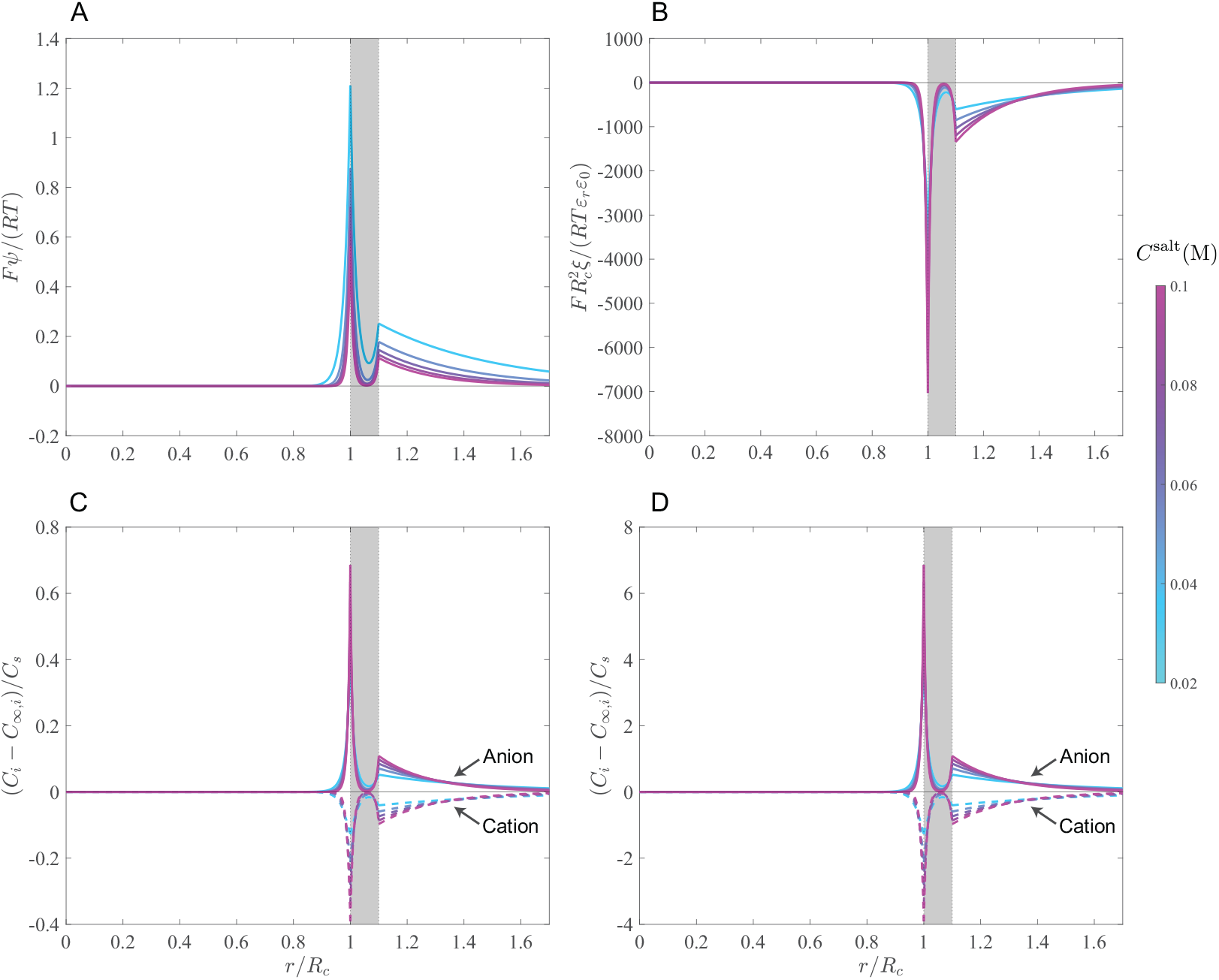
Steady-state solutions of species mass-balance and Maxwell’s first equations at *R*_*c*_ = 10^−7^ m, *σ* = 0.01 C/m^2^, and *σ*_*r*_ = 0.2. (A) Electric potential, (B) volume charge density, (C) concentration of cations and anions associated with salt-I, and (D) concentration of cations and anions associated with salt-II. Here, *C*_*∞,i*_ denotes the farfield concentration of ion *i* in the ocean, arising from dissociation of the respect salt in water. Shaded areas indicate the position of the membrane along the *r*-axis. The scale on the *r*-axis, where the ocean lies, is stretched by a factor 20 to better show radial profiles.

### Trade-off Between Stability and Sensitivity to Surface Charge Could Have Driven Early Evolution of Membrane Potential

The steady-state results presented in the last section indicated that large positive membrane potentials are achievable if mineral surfaces can attain large surface charge densities. Of course, how large a surface charge density a mineral can attain is dictated by the laws of thermodynamics and depends on several factors, such as the temperature, pressure, and ionic composition of the solution it is exposed to [17]. To better understand if other constraints could have restricted the range of achievable membrane potentials, we systematically studied the steady-state solutions of Eqs. (1) and (2) as a function of surface charge density *σ*. We constructed parametric steady state solutions with respect to *σ* at fixed *R*_*c*_, *C*^salt^, and *σ*_*r*_ using branch continuation methods [18] (see “SI: Constructing Steady-State Solution Branches”) and determined stability along the solution branches using linear stability analysis (see “SI: Stability of Steady-State Solutions”).

Steady-state solution branches, represented in scaled membrane potential 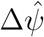 versus scaled surface charge density 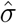 diagrams (see “SI: Nondimensionalization of Governing Equations” for definition of scaled quantities), exhibit a distinct trend at large (Fig. 3A) and small (Fig. 3B) cell radii. In general, 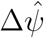 increases linearly with 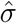 at small 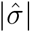, asymptotically plateauing at large 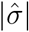. Here, 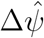 increases with 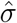 at a faster rate for smaller cells. However, when solution branches are represented with respect to dimensional quantities, the opposite trend is observed; that is, Δ*ψ* increases with *σ* at a faster rate for larger cells. Moreover, the linear stability analysis shows that, in general, there is a neighborhood around 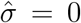, in which steady state solutions are stable. Increasing 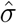 beyond a critical value results in stability loss. Therefore, unboundedly large membrane potentials cannot be achieved by arbitrarily increasing the membrane surface charge density, even if it was allowed by the laws of thermodynamics.

**Figure 3:**
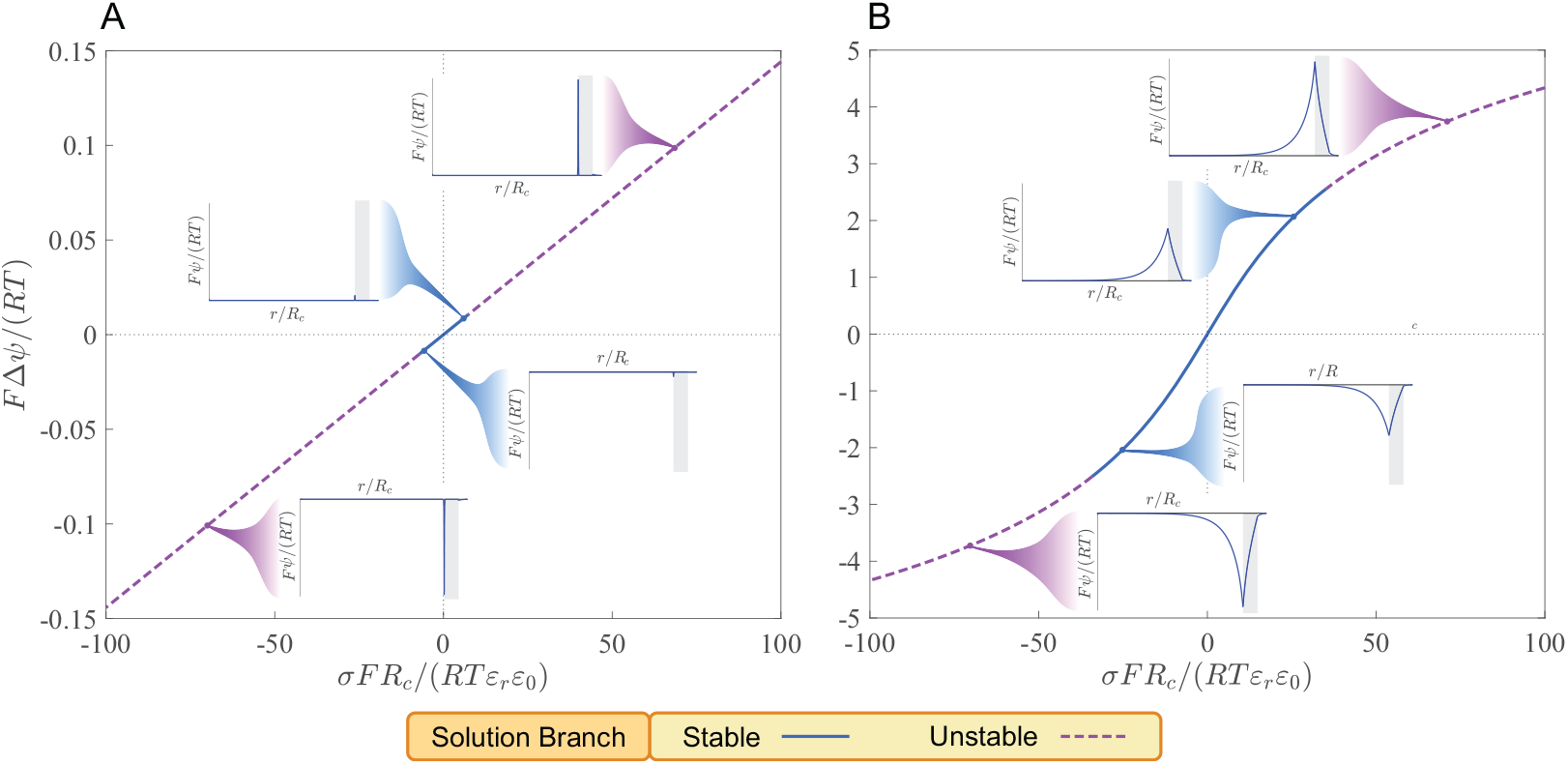
Steady-state solutions of the membrane potential Δ*ψ* parametrically represented with respect to the surface charge density *σ* at *C*_*∞*_ = 0.2 M and *σ*_*r*_ = 0.02 with (A) *R*_*c*_ = 10^−6^ m and (B) *R*_*c*_ = 10^−8^ m. Inset figures show radial profile of the electric potential *ψ* at selected points along the steady-state solution branches. Shaded areas indicate the position of the membrane along the *r*-axis. Stability analysis was performed for *ϑ* = 0.1. The tortuosity coefficient *ϑ* only affects stability without altering steady-state solutions.

Other parameters besides *σ* could have controlled the development of the membrane potential in primitive cells by altering the steady states and their stability, such as the total salt concentration in the ocean *C*^salt^ and the tortuosity coefficient of the membrane *ϑ*. These characterize the ionic composition of the ocean and microstructural properties of the membrane, respectively. The tortuosity coefficient is defined as the ratio of the effective diffusivity in the membrane and bulk diffusivity for any given ion [19, 20] and measures how much the tortuous microstructure of porous membranes hinders diffusive transport. It only affects the stability of ion distributions in our protocell model, but not the steady-state solutions (see “SI: Stability of Steady-State Solutions”).

We studied how *C*^salt^ could influence steady-state solution branches in 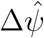 versus 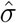 diagrams at large (Figs. S2A,C) and small (Figs. S2B,D) cell radii by constructing solution branches at fixed values of *C*^salt^. We then determined stability along solution branches for *ϑ* = 0.05 (Figs. S2A,B) and *ϑ* = 0.1 (Figs. S2C,D). In general, lowering *C*^salt^ increases the sensitivity of 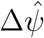 to 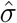. It also reduces the range of 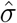, in which steady-state solutions are stable. Lowering *ϑ* has a similar destabilizing effect by reducing the stable range of 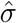. Interestingly, the stable regions of *σ* determined in all these case studies are within the experimentally measured ranges of surface charge densities at mineral-water interfaces (see “SI: Surface Charge of Minerals”).

Overall, we observe a trade-off between stability and the sensitivity of Δ*ψ* to *σ*, manifesting itself in two distinct ways (Fig. 4). The first is connected with the size of the cell. Here, Δ*ψ* is more sensitive to *σ* at larger cell radii. However, the range of *σ*, in which stable ion distributions can be achieved becomes more restricted in return. Restrictions on *σ*, in turn, place a constraint on the maximum achievable Δ*ψ*. The second is related to the composition of the ocean. In this case, a lower *C*^salt^ provides Δ*ψ* that is more sensitive to *σ* at the expense of destabilization of ion distributions.

**Figure 4:**
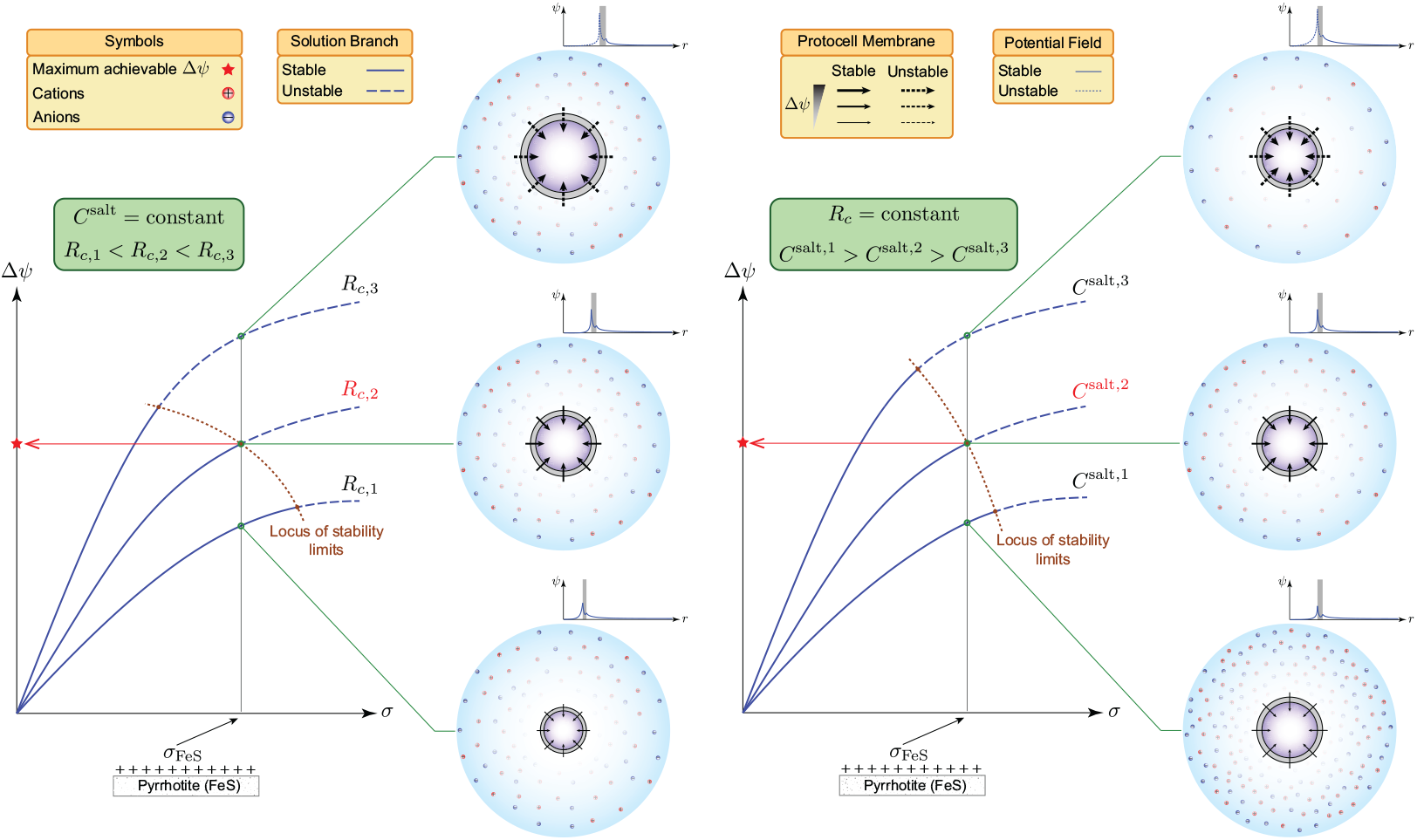
Trade-off between stability and the sensitivity of the membrane potential Δ*ψ* to surface charge density *σ*. Solution branches for a given tortuosity coefficient *ϑ* are qualitatively plotted for several *R*_*c*_ at fixed *C*^salt^ (left diagram) and several *C*^salt^ at fixed *R*_*c*_ (right diagram). Quantitative solution branches and their stability limits are shown in Figs. S2 and S3, respectively. In both cases, there is a critical *σ* and Δ*ψ*, beyond which stability is lost. At fixed *C*^salt^ (left diagram), larger membrane potentials can be achieved for a given *σ* when *R*_*c*_ is larger, although stability is lost at a smaller *σ*. Similarly, at fixed *R*_*c*_ (right diagram), larger membrane potentials can be realized for a given *σ* when *C*^salt^ is smaller, but stability is lost at a smaller *σ*. Consequently, for a given *σ* corresponding to the constituents of the membrane (*e.g.*, FeS), there is an intermediate *R*_*c*_ at fixed *C*^salt^ (highlighted in red in the left diagram) or an intermediate *C*^salt^ at fixed *R*_*c*_ (highlighted in red in the right diagram) that furnishes the maximum achievable Δ*ψ*. Arrows on the membrane point in the direction, in which Δ*ψ* exerts force on negative ions.

### Electroneutrality Provides Selective Advantage by Minimizing Osmotic Pressure and Concentration Heterogeneity

So far, we focused our discussion on the role that the membrane potential could have played as a barrier, preventing organic molecules, synthesized by earliest metabolic reactions, from dissipating into the ocean. In modern cells, this role has been taken over by lipid bilayer membranes, while the membrane potential together with membrane proteins are involved in membrane bioenergetics [21], pH and metal-ion homeostasis [11], controlling material flow into and out of the cell [22, 12], and stress regulation [23, 10]. As previously stated, a nonzero membrane potential in protocells with ionpermeable membranes would have necessitated the violation of electroneutrality. By contrast, electroneutrality is always maintained in modern cells by controlling ion transfer across the membrane using sophisticated regulatory networks [24, 25]. It may be plausible to surmise that lipid membranes were assimilated by primitive cells from early stages [26] once terpenoid synthesis pathways were incorporated into primordial reaction networks [5]. However, what selective pressures could have driven the evolution of charge distributions towards electroneutrality by selecting for ion-impermeable boundary structures? In the following, we highlight two reasons for why electroneutrality could have been selected for at early stages of evolution, namely (i) the osmotic crisis and (ii) concentration heterogeneity.

The osmotic crisis refers to membrane breakup as a result of ion accumulation inside the cell [22]. This universal phenomenon, which applies to primitive and modern cells, occurs due to the uptake of charged cofactors, reducing agents, carbon sources, and energy sources. The elevated ion concentrations in the cell lower the water activity compared to the surrounding water [27]. Consequently, water is driven through the membrane into the cell, causing the membrane to break. To better understand how electroneutrality could have contributed to this process, we considered a simplified case, involving an electrolyte solution with an anion and a cation (see “SI: Electroneutrality and Mechanical Stability of Protocells”). We used Pitzer’s model [28] to estimate the osmotic coefficient of the solution. Given that the osmotic coefficient is mainly a function of the ionic strength [29], we sought to determine how the osmotic coefficient varies with the net charge of the solution if the ionic strength is maintained constant. We found that, at fixed ionic strength, of all possible ionic compositions, the one yielding an electroneutral solution minimizes the osmotic coefficient. This implies that, primordial reaction networks could have progressed for longer times under electroneutral conditions by taking up all the necessary ingredients from the surroundings without undergoing any catastrophic event. As a result, the likelihood of early metabolism incorporating additional steps to synthesize biomolecules of higher complexity could have improved (Fig. 5).

**Figure 5:**
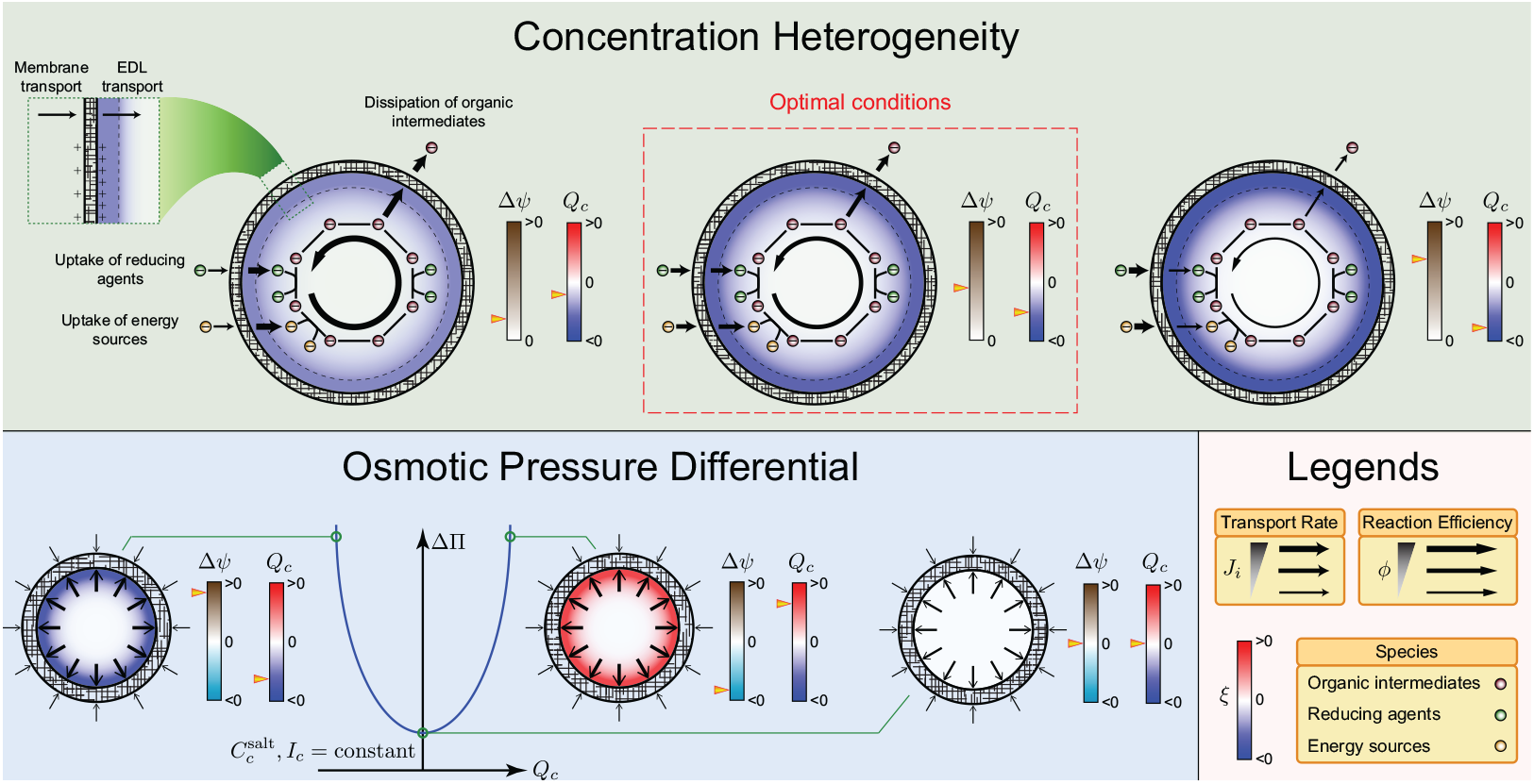
Selective advantages provided by electroneutrality. Selective pressures are examined in a protocell model (Fig. 1), in which primordial reactions occur. Large positive membrane potentials Δ*ψ* increase the transport rate of negatively charged reducing agents and energy sources, while attenuating the dissipation of the organic intermediates of metabolic reactions. They also induce a nonuniform positive electric potential field, elevating the concentration of anions in the cell. The resulting negative charge-density distribution (*ξ*(*r*) < 0) leads to heterogeneous concentration distributions for charged reactive species, which, in turn, diminish reaction efficiencies. Small membrane potentials do not affect the reaction efficiencies significantly. However, they can not alleviate the dissipation of the organic intermediates as effective either. Therefore, parameter values furnishing an intermediate range of positive membrane potentials would have been optimal for the operation of early metabolic cycles. Moreover, if the total salt concentration 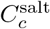 and ionic strength *I*_*c*_ in the cell are held constant, then violation of electroneutrality (|*Q*_*c*_| > 0 with *Q*_*c*_ the total charge in the cell) increases the osmotic pressure differential ΔΠ across the membrane. Hence, electroneutrality could have minimized catastrophic events in primitive cells due to osmotic crisis, promoting the evolution of complex metabolic networks by providing structural stability.

Electroneutrality could also have enhanced the efficiency of early metabolic reactions by creating homogeneous concentration distributions inside primitive cells. To clarify this point, consider a scenario, where positively charged membrane surfaces have generated a positive membrane potential in a primitive cell. The positive membrane potential drives negative inorganic ions into the cell, creating a negatively charged shell (*i.e.*, electric double layer) next to the inner surface of the membrane. Next, suppose that a negatively charged organic molecule is consumed by some of the metabolic reactions occurring in the cell. To efficiently utilize this molecule, the cell must maximize its concentration by minimizing its rate of transport into the ocean. This minimization is accomplished by the positive membrane potential. However, the concentration of the molecule is only increased at the inner surface of the membrane since it cannot readily diffuse past the charged shell to mix and react with other molecules. Therefore, organic molecules are nonuniformly distributed inside the cell, diminishing mixing and reaction efficiencies (Fig. 5).

To quantify the extent to which nonuniform concentration distributions—referred to as concentration heterogeneity—can reduce apparent reaction rates, we solved the species mass-balance equation Eq. (2) for a reactive species *B* consumed according to the first-order rate law *r*_*B*_ = −*kC*_*B*_ in a protocell with a prescribed volume-charge-density distribution (see “SI: Concentration Heterogeneity and Reaction Efficiency”) to ascertain the radial concentration distribution *C*_*B*_(*r*) and its volume average 〈*C*_*B*_〉. We then used the integral form of Eq. (2) to relate the total uptake flux of *B* and apparent reaction rate 〈*r*_*B*_〉 ≔ ∫_*V*_*r*_*B*_d*V/V* = −*k*〈*C*_*B*_〉. Given the linear proportionality between 〈*r*_*B*_〉 and 〈*C*_*B*_〉 in this case, 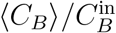 may be regarded as a measure of how much concentration heterogeneity can diminish or enhance apparent reaction rates (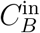 denotes the concentration of *B* at the inner surface of the membrane). Lastly, we examined how 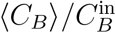 is affected by the rate constant and surface charge density (Fig. S4).

Our analysis revealed two distinct ways, in which concentration heterogeneity can affect apparent reaction rates. The first is due to the local consumption of *B*, always reducing the apparent reaction rate of *B*, irrespective of the background charge distribution arising from the inorganic ions. The second is related to the charged shell discussed above. If the shell and *B* have opposite charges, the apparent reaction rate is enhanced (Fig. S4, dashed lines), and if they have like charges, the apparent reaction rate is reduced (Fig. S4, solid lines). As it relates to the origin of metabolism, positive membrane potentials could have favored the evolution of early metabolism by minimizing the dissipation of organic intermediates into the ocean. However, they could also have degraded mixing and reaction efficiencies, thereby driving the evolution of ion-impermeable membranes, specialized ion channels, and active transport systems to maintain an electroneural intracellular environment and minimize the interaction of membrane transport and electric double layers.

## Discussion

The plausibility of life originating from prebiotic metabolic cycles promoted by naturally occurring catalysts on the primitive Earth (*i.e.*, the metabolism-first hypothesis) has been subject to much scrutiny [30, 31, 15]. Perhaps one of the main criticisms of this hypothesis is the supposed improbability of metabolic cycles, comprising several distinct steps, that could have been sustained stably by prebiotic catalysts [30, 31]. From this standpoint, it is deemed unlikely that any assortment of minerals on the primitive Earth could have been efficient and specific enough to catalyze diverse sets of metabolic reactions to synthesize the essential building blocks of life [31]. These are organic molecules that must have been produced in high enough concentrations to support the subsequent evolution of early metabolic cycles towards networks of higher complexity—a challenging task to accomplish without enzymes, given the comparatively poor efficiency of inorganic catalysts [8].

We addressed the foregoing criticism by proposing a mechanism, through which the concentration of organic intermediates of early metabolic cycles could have been enhanced without sophisticated macromolecular structures or polymerization machinery, which are believed to have been later products of evolution [32]. In this mechanism, membrane surfaces in primitive cells are assumed to have been positively charged due to the accumulation of transition metals. Then, these charged surfaces could have induced a positive membrane potential, which, in turn, would have concentrated the organic intermediates of early metabolic cycles inside primitive cells. We demonstrated the feasibility of this mechanism by developing a protocell model and quantitatively estimating achievable membrane potentials from first principles by solving Maxwell’s first law and mass-balance equations. We showed that positive membrane potentials comparable in magnitude to those observed in modern bacteria could have been generated in primitive cells for typical charge densities arising from transition-metal surfaces.

To better understand how a positive membrane potential could have developed, we constructed the steady-state solutions of Maxwell’s first law and mass-balance equations. We found that positive membrane surface charges could induce a nontrivial electric potential field and a positive membrane potential. The resulting membrane potential is proportional to the surface charge density and inversely proportional to the total concentration of nonreactive ions in the primitive ocean. Furthermore, our numerical experiments indicated that violation of electroneutrality inside the cell and membrane is essential to generate a nonzero membrane potential. However, the steady-state results alone did not place any upper limit on the maximum achievable membrane potential.

Thus, we examined the stability of the steady-state solutions using linear stability analysis to identify possible constraints that could have restricted the magnitude of the membrane potential in primitive cells. Our results suggested that, for any given ionic composition of the primitive ocean, there is a critical surface charge density and membrane potential, beyond which concentration distributions in the cell and membrane are unstable. Moreover, we found that there is a trade-off between this stability bound and the sensitivity of the membrane potential to surface charge density: Parameter values leading to higher sensitivities result in a smaller range of surface charge densities, for which concentration distributions are stable. Beside destabilization, large surface charge densities could also have induced heterogeneous concentration distributions for organic molecules, cofactors, and energy sources inside primitive cells, adversely affecting the reaction efficiencies of early metabolic cycles. This is yet another reason for why arbitrarily large membrane potentials could not have been achieved.

Lastly, our quantitative analysis revealed that the conditions on the primitive Earth could have been primed for the emergence of first metabolic cycles, perhaps more than previously thought. The feasibility of these cycles in our model relies on the existence of a positive membrane potential. In fact, the concordance between the interconnection of metabolic reactions and the membrane potential in primitive and modern cells is an important feature of our protocell model. It implies that, the operation of the membrane potential and metabolism were deeply intertwined from the outset and continued to persist throughout the evolutionary history of life. Furthermore, our results suggested that, sufficiently large membrane potentials could have been realized for intermediate ranges of surface charge density, cell size, and ion concentrations in the ocean to support the evolution of stable and self-sustaining metabolic cycles. These ranges may be regarded as constraints exerting selective pressure on the evolution of early metabolism. They would likely have been more restrictive at the beginning and were relaxed once lipid membranes and specialized ion channels had emerged, which, in turn, would have rendered primitive cells more robust to environmental uncertainties.

Overall, this study provides a strong impetus for further rigorous and quantitative investigations into mechanistic models of first metabolic cycles and their early evolutionary stages, elucidating their transition into self-sustaining and complex biochemical networks. More broadly, our results suggest that the strong interconnections between several cellular processes (*e.g.*, controlled membrane potential and membrane transport, charge balance, ion homeostasis, metabolism) were as essential to primitive cells as they are to extant life. These are fundamental processes that shape many phenotypic characteristics of modern cells, possibly more than currently understood. Therefore, we expect that these fundamental processes will be formalized more systematically in the future for biological systems and incorporated into realistic single-cell models to emerge in the coming decade.

## Supplementary Information

### Protocell Model

We first describe the protocell model discussed in the main text that we proposed to study the origin of the membrane potential and electroneutrality. The model comprises three regions: (i) Cell, (ii) membrane, and (iii) ocean (Fig. S1). The cell is a sphere of radius *R*_*c*_ enclosed by a porous membrane of thickness *d*, lying at the bottom of the primitive ocean (Fig. 1A). Our goal is to determine the conditions, under which a positive membrane potential can develop across the membrane. However, we consider a more general case, where positive and negative membrane potentials can be induced by positively and negatively charged surfaces of the membrane. The inner and outer surfaces of the membrane are assumed to have the same charge with the respective surface charge densities *σ*^in^ and *σ*^out^. However, the magnitude of the surface charge density on the inner surface is assumed to be always greater than on the inner surface (Fig. 1A). These positively and negatively charged surfaces could have arisen from accumulation of transition-metal and clay minerals, respectively [5]. In our model, the inner and outer surface charge densities are specified using two parameters according to *σ*^in^ = *σ* and *σ*^out^ = *σ*_*r*_*σ*, where the the surface charge density *σ* and surface charge density ratio *σ*_*r*_ are given parameters. Other fixed parameters of the model that were used to generate the plots in this document and the main text are summarized in Table S1.

The ocean in our model is assumed to be electroneutral. We further assume that the ionic composition of the ocean arises from a complete dissociation of monovalent salts. For simplicity, we only consider two monovalent salts, we refer to as salt-I and salt-II, which yield equal amounts of the respective cation and anion in the ocean upon dissociation in water. Let 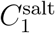 and 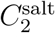 denote the concentration of salt-I and salt-II, respectively. Then, the total salt concentration 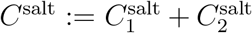 is half the total ion concentration *C*_*∞*_ in the ocean due to a complete salt dissociation. Therefore, when *C*^salt^ and 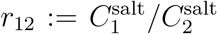 are given, the ionic composition of the ocean is fully specified.

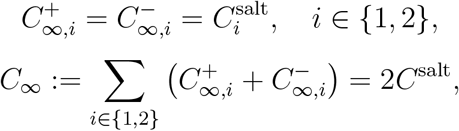

where 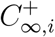 and 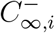 are the concentration of the cation and anion arising from salt *i*. We generally denote the concentration of ions (cations or anions) by *C*_*∞,i*_ without referring to the index of salt, where *i* here is the index of ions in the system. The composition of the ocean is then imposed as far-field boundary conditions to solve Maxwell’s first law and species mass-balance equations in the three computational domains shown in Fig. S1.

**Figure S1:**
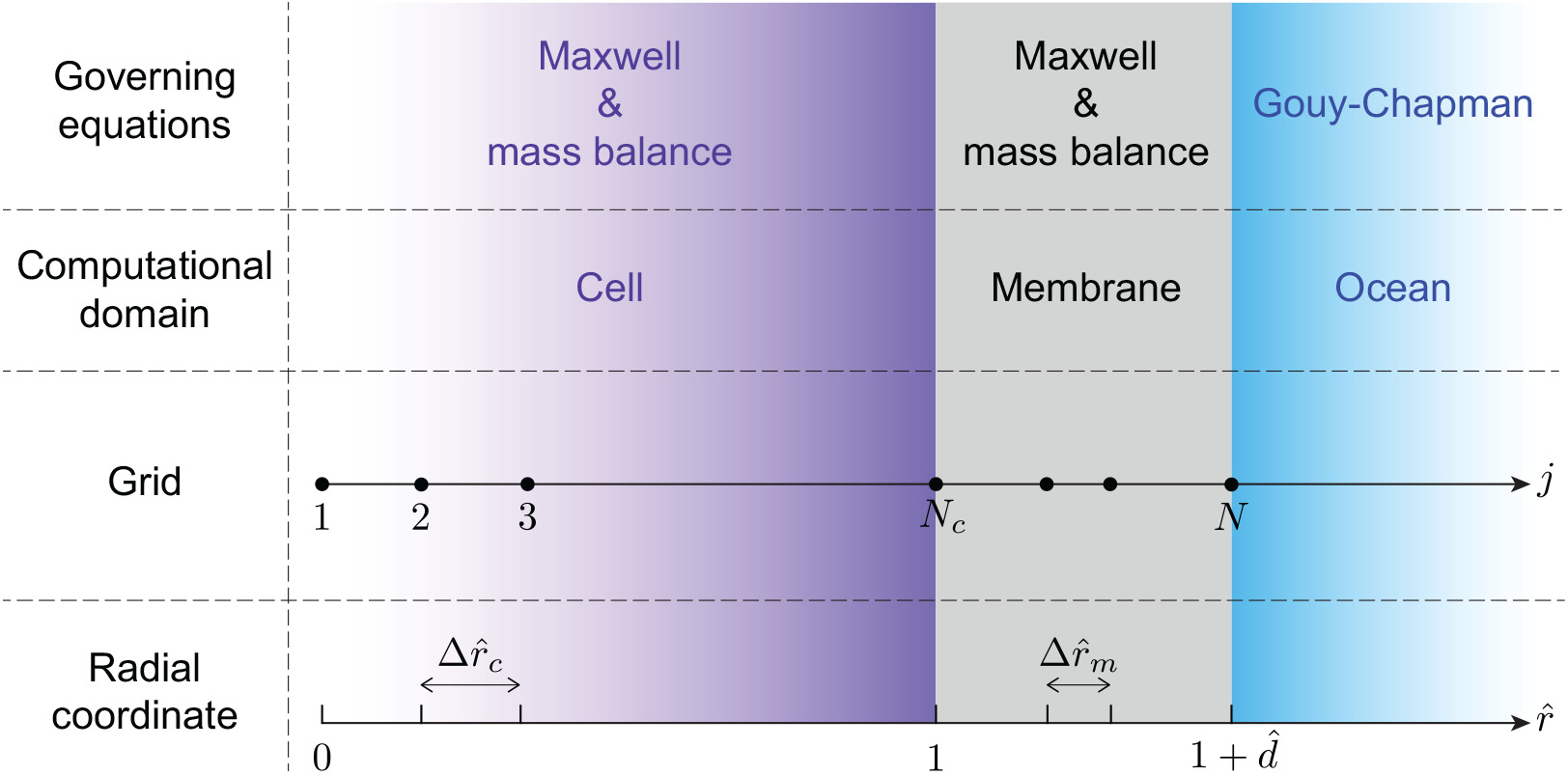
Computational domains in the protocell model of life’s origins described in Fig. 1 and the grid used to discretize the governing equations. The model comprises three computational domains, namely the cell, membrane, and ocean. Maxwell’s first law and species mass-balance equations are solved in the cell and membrane to ascertain the electric potential field and concentration distributions. However, the surface potential on the outer surface of the membrane, electric potential field, and concentration distributions in the ocean are approximated by the Gouy-Chapman theory [33, Section 5.3].

### Governing Equations

To ascertain the membrane potential for any given set of model parameters, we compute the electric potential field in the foregoing three computational domains by solving Maxwell’s first law

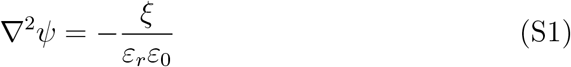

and species mass-balance equations

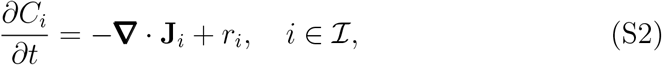

where *ψ* is the electric potential, *ξ* volume charge density, *ε*_*r*_ relative permittivity of water, *ε*_0_ vacuum permittivity, 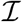 the index set of all the species involved in the system with *C*_*i*_, **J**_*i*_, and *r*_*i*_ the concentration, flux vector, and production rate of species *i*. The three computational domains, in which to solve Eqs. (S1) and (S2) are represented as

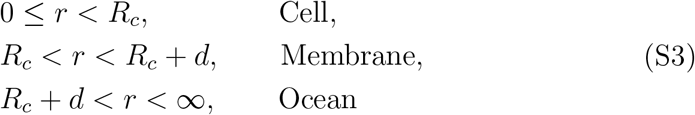

with the boundary conditions

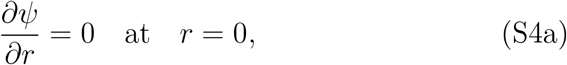

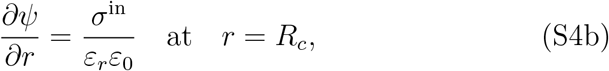

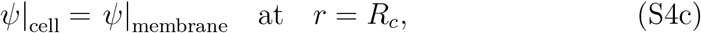

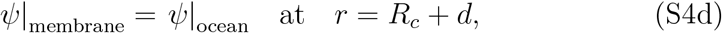

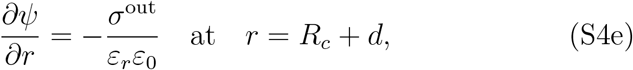

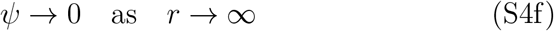

for the electric potential and

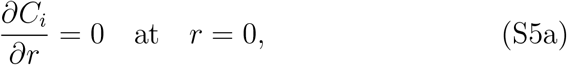

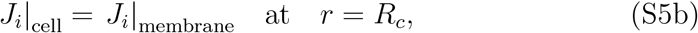

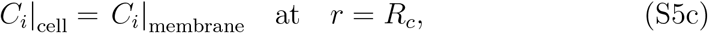

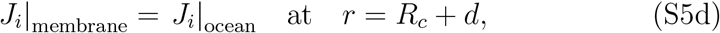

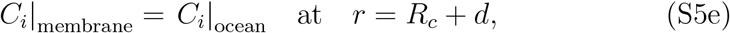

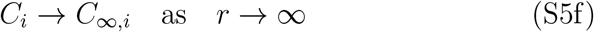

for the species concentrations, where *J*_*i*_ ≔ **J**_*i*_ · **e**_*r*_ with **e**_*r*_ the unit vector along the *r*-axis in the spherical coordinate system. The boundary conditions Eqs. (S4b) and (S4e) arise from a charge-balance constraint applied in the electric double layer theory [33, Section 5.3]. It requires that the total charge resulting for the accumulation of the counter ions in the domain exposed to a charged surface counterbalances the total charge of the surface. Applying this constraint to the electric double layers formed in the cell and ocean yields

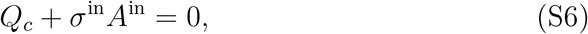

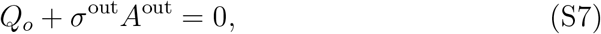

where *Q*_*c*_ and *Q*_*o*_ are the total charges accumulated in the cell and ocean with *A*^in^ and *A*^out^ the areas of the inner and out membrane surfaces. The total charges can then be related to 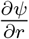 using Gauss’s law, which is the integral form of Maxwell’s first law. For example, Eq. (S4b) can be derived from Eq. (S6) in the following way

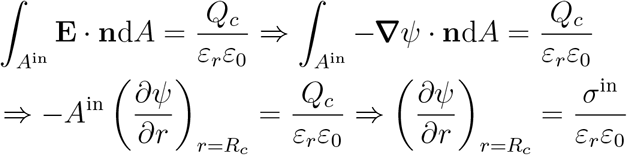

with **E** the electric field and **n** unit outward normal vector to *A*^in^. Equation (S4c) can be similarly derived from Eq. (S7).

**Table S1:**
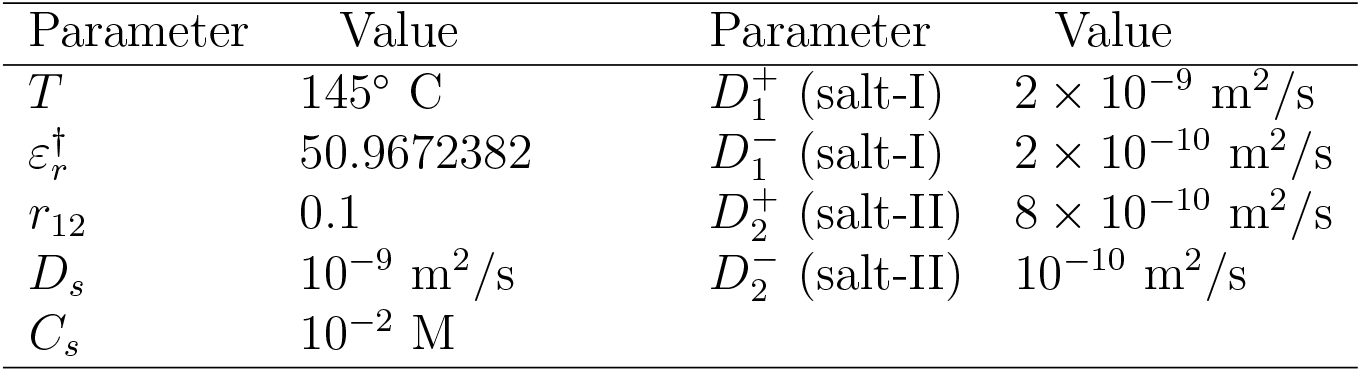
Parameters used for all the case studies presented in the main text and supplementary information.

| Parameter | Value | Parameter | Value |
| --- | --- | --- | --- |
| $T$ | 145° C | $D_1^+$ (salt-I) | $2 \times 10^{-9} \text{ m}^2/\text{s}$ |
| $\varepsilon_r^\dagger$ | 50.9672382 | $D_1^-$ (salt-I) | $2 \times 10^{-10} \text{ m}^2/\text{s}$ |
| $r_{12}$ | 0.1 | $D_2^+$ (salt-II) | $8 \times 10^{-10} \text{ m}^2/\text{s}$ |
| $D_s$ | $10^{-9} \text{ m}^2/\text{s}$ | $D_2^-$ (salt-II) | $10^{-10} \text{ m}^2/\text{s}$ |
| $C_s$ | $10^{-2} \text{ M}$ | | |

<sup>†</sup>Estimated using the revised Helgeson-Kirkham-Flowers equation of state at 100 bar and in the temperature range 120–145° [34].

The volume charge density in Eq. (S1) is determined from the concentration of the species in the system

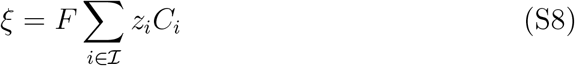

with *F* the Faraday constant and *z*_*i*_ the valence of species *i*. The flux vector consists of a diffusive and an electric-potential component, which is expressed

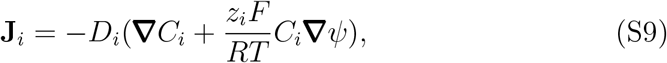

where *R* is the universal gas constant and *T* temperature.

Before proceeding to the steady-state solutions of Eqs. (S1) and (S2), it is helpful to examine these equations separately for two groups of species. Here, we classify the species involved in our model into a reactive and non-reative group. The reactive species are those that would have participated in early metabolic reactions occurring inside the cell, such as reducing agents, energy sources, and organic molecules. The nonreactive species are the inorganic ions, which would have been present in the primitive ocean. The rates of nonenzymatic reactions in the earliest metabolic cycles would have been much smaller than enzymatic reactions in modern metabolic networks. Therefore, the concentrations of nonreactive species involved in primitive reactions would have been much smaller than those of metabolites in modern organisms and especially of inorganic ions in the primitive ocean [14]. Accordingly, the volume charge density in Eq. (S1) can be approximated

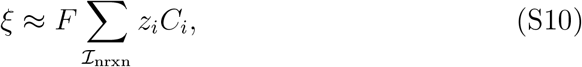

where 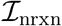 the index set of nonreactive species. Using this approximation along with *r*_*i*_ = 0 for nonreactive species, one can solve Eqs. (S1) and (S2) for nonreactive species independently of the reactive species. Once the electric potential has been ascertained in this manner, the species mass-balance equations for reactive species can be solved without needing to couple them to Maxwell’s first law.

We conclude this section by highlighting the main assumptions used in the remainder of this document to simplify the governing equations and their solutions:

- Governing equations (steady states and transient perturbations) inherit spherical symmetry from the spherical geometry of the protocell model.
- Surface charge density can vary independently of other model parameters, such as cell radius, membrane thickness, and the ionic composition of the ocean.

The second assumption is only relevant to parametric studies of the steady-state solutions with respect to the surface charge density *σ*. As will be discussed later (see “Constructing Steady-State Solution Branches”), we construct steady-state solution branches with respect to *σ* at fixed *C*^salt^. This is, of course, a simplifying assumption because when the thermodynamic state of the system is specified (*i.e.*, when the temperature, pressure, and ionic composition are given), *σ* is determined by the thermodynamic constraints arising from the equilibrium of the charged surface and the electrolyte solution it is subject to [17]. Therefore, in general, *σ* and *C*_*∞*_ cannot vary independently of one another.

### Nondimensionalization of Governing Equations

To alleviate computational errors associated with the scaling of the protocell model that arise from the numerical solutions of the governing equations, we introduce the dimensionless quantities

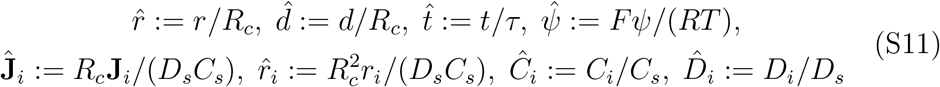

to nondimensionalize these equations, where *C*_*s*_ and *D*_*s*_ are concentration and diffusivity scales. Accordingly, after applying the spherical symmetry assumption, the dimensionless forms of Eqs. (S1) and (S2) are obtained

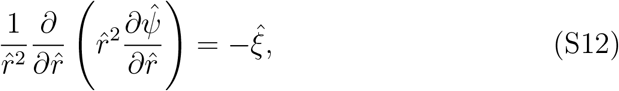

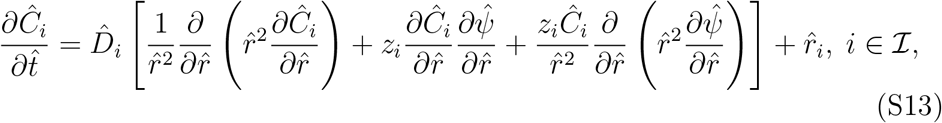

which are to be solved subject to

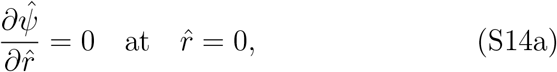

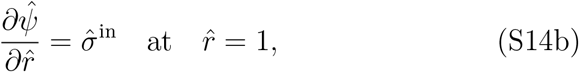

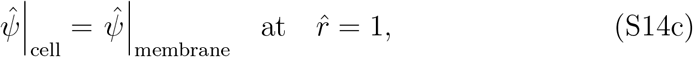

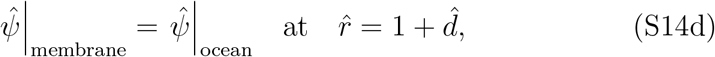

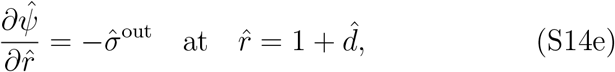

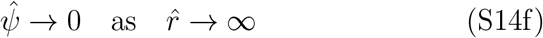

for the electric potential and

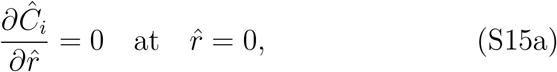

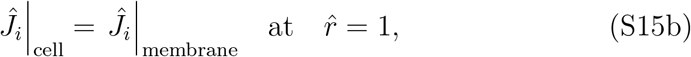

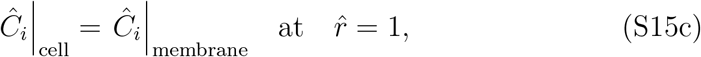

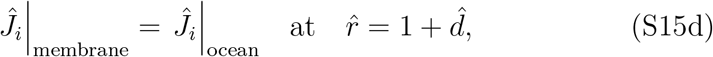

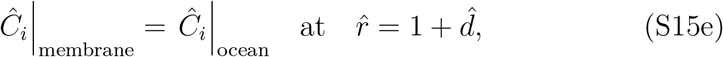

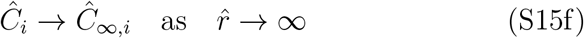

for the species concentrations, where 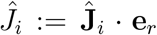. Several dimensionless parameters appear in these equations, the definitions of which are

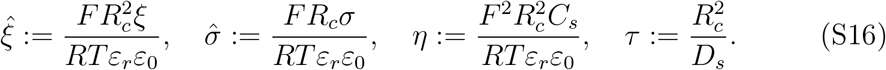

Note that, throughout this document, the dimensionless forms of all the other concentrations, electric potentials, and surface charge densities are denoted as the corresponding hatted quantities and defined similarly. Using these dimensionless parameters, Eq. (S9) is nondimensionalized as

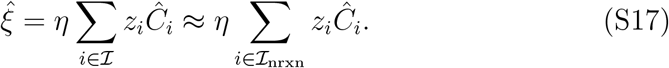

### Steady State Solutions

In this section, we present the steady state solutions of Eqs. (S12) and (S13) for nonreactive species. We approximate the steady-state solutions in the ocean using the Gouy-Chapman theory for simplicity [33] (see “Electric Potential Field in Ocean from Gouy-Chapman Theory”) and compute numerically exact solutions of Maxwell’s first law and species mass-balance equations in the cell and membrane. Because there are no sources or sinks for nonreactive species in the cell or membrane, 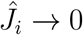 for 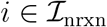 as the solutions of Eqs. (S12) and (S13) approach their steady states. This observation allows to simplify the construction of steady-state solutions as demonstrated in the following

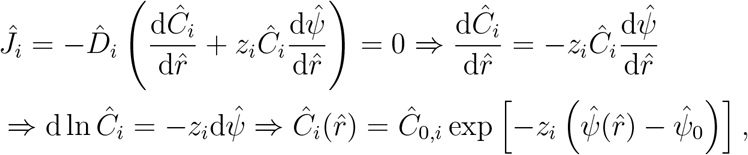

where 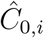 and 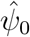 are the concentration of species *i* and electric potential on one of the boundaries of the computational domain. Note that this simplification does not apply to reactive species, for which 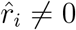, because steady-state fluxes can generally be nonzeros.

For the cell and membrane, the boundary of interest is *A*^in^ and *A*^out^, respectively. The surface concentrations and surface potential on *A*^out^, which are ascertained from the Gouy-Chapman theory in the ocean provide the boundary conditions for the membrane. The surface concentrations and surface potential on *A*^in^ from the solution of the membrane, in turn, furnish the boundary conditions for the cell. Once these boundary conditions are substituted in the general expression derived above, the following concentration distributions in the cell and membrane are obtained

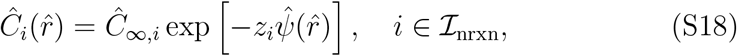

which describe the functional dependence of 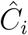 on 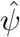 both in the cell and membrane. However, the concentration distributions 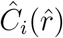 in the cell and membrane are not the same because the electric potential field 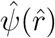 in these domain are different. Substituting Eq. (S18) in Eq. (S12) using Eq. (S17) furnishes

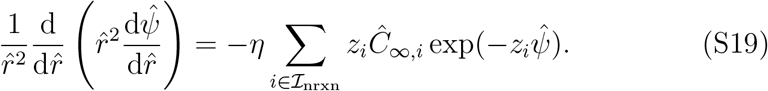

This form of Maxwell’s first law needs not be coupled to the species mass-balance equations Eq. (S13) to provide the electric potential field. We numerically solve this equation using finite-difference methods (see “Numerical Approximation of Steady-State Solutions”) subject to the boundary conditions Eqs. (S14a)–(S14d) to compute 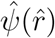 in the cell and membrane for a given set of model parameters. Once the electric potential field has been computed, it can be back-substituted in Eq. (S18) to provide the concentration distributions.

### Electric Potential Field in Ocean from Gouy-Chapman Theory

As previously stated, we approximate the electric potential field and concentration distribution of ions in the ocean using the Gouy-Chapman theory. This theory provides analytical solutions for 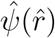 and 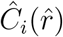 when only monovalent ions are present in an electrolyte solution and the domain is one-dimensional in the Cartesian coordinate system [33, Section 5.3]. In this theory, ion concentrations are explicitly expressed as functions of the electric potential field. The functional form of these expressions is identical to that in Eq. (S18). The electric potential field and surface potential are given by

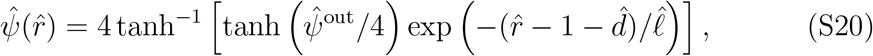

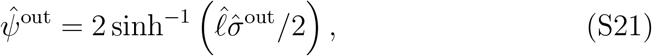

where 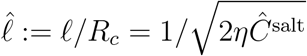 with *ℓ* the Debye length defined as

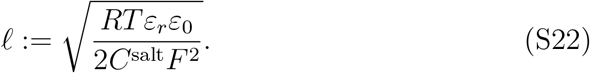

### Numerical Approximation of Steady-State Solutions

We briefly discuss the numerical techniques, with which to solve Eq. (S19). We discretize Eq. (S19) over the grid shown in Fig. S1 and approximate the first and second derivatives of *ψ* that arise from its left-hand side using fourth-order finite-difference schemes (see Tables S2 and S3). Substituting these approximations in Eq. (S19) yields a nonlinear system of equations

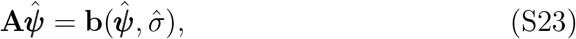

where 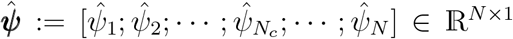 is the vector of the electric potentials evaluated at the nodes of the grid shown in Fig. S1. Here, **A** ∈ ℝ^*N×N*^ is a constant matrix comprising the coefficients of the discretization schemes and **b** ∈ ℝ^*N×*1^ is a variable vector and nonlinear in 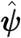. It results from the right-hand side of Eq. (S19) evaluated at the grid points and the boundary conditions Eqs. (S14a)–(S14d).

**Table S2:** Forth-order finite-difference schemes to approximate the first derivative of a function 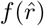 on the grid shown in Fig. S1. Expressions in the Scheme column are derivatives evaluated at 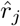, where 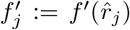. Note that 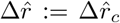 when 1 ≤ *j* ≤ *N*_*c*_ and 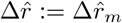 when *N*_*c*_ + 1 ≤ *j* ≤ *N* (see Fig. S1).

| Index | Scheme |
| --- | --- |
| $j = 1$ | $\frac{-25f_j + 48f_{j+1} - 36f_{j+2} + 16f_{j+3} - 3f_{j+4}}{12\Delta\hat{r}}$ |
| $j = 2, N_c + 1$ | $\frac{-3f_{j-1} - 10f_j + 18f_{j+1} - 6f_{j+2} + f_{j+3}}{12\Delta\hat{r}}$ |
| $\begin{cases} 3 \leq j \leq N_c - 2 \\ N_c + 2 \leq j \leq N - 2 \end{cases}$ | $\frac{f_{j-2} - 8f_{j-1} + 8f_{j+1} - f_{j+2}}{12\Delta\hat{r}}$ |
| $j = N_c - 1, N - 1$ | $\frac{-f_{j-3} + 6f_{j-2} - 18f_{j-1} + 10f_j + 3f_{j+1}}{12\Delta\hat{r}}$ |
| $j = N_c, N$ | $\frac{3f_{j-4} - 16f_{j-3} + 36f_{j-2} - 48f_{j-1} + 25f_j}{12\Delta\hat{r}}$ |

**Table S3:** Forth-order finite-difference schemes to approximate the second derivative of a function 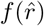 on the grid shown in Fig. S1. Expressions in the Scheme column are derivatives evaluated at 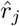, where 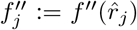. Note that 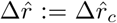 when 1 ≤ *j* ≤ *N*_*c*_ and 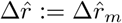 when *N*_*c*_ + 1 ≤ *j* ≤ *N* (see Fig. S1).

| Index | Scheme |
| --- | --- |
| $j = 2, N_c + 1$ | $\frac{10f_{j-1} - 15f_j - 4f_{j+1} + 14f_{j+2} - 6f_{j+3} + f_{j+4}}{12\Delta\hat{r}^2}$ |
| $\begin{cases} 3 \leq j \leq N_c - 2 \\ N_c + 2 \leq j \leq N - 2 \end{cases}$ | $\frac{-f_{j-2} + 16f_{j-1} - 30f_j + 16f_{j+1} - f_{j+2}}{12\Delta\hat{r}^2}$ |
| $j = N_c - 1, N - 1$ | $\frac{f_{j-4} - 6f_{j-3} + 14f_{j-2} - 4f_{j-1} - 15f_j + 10f_{j+1}}{12\Delta\hat{r}^2}$ |

We solve Eq. (S23) iteratively in two steps:

- Pseudo-linear step: The procedure starts from a crude initial guess 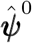. At iteration *n*, 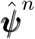 is computed by solving the linearized system 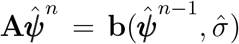, where 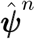 and 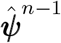 denote the vector of electric potentials at iterations *n* and *n*−1, respectively. This step is relatively robust with respect to the initial guess but converges slowly to steady-state solutions. The approximate solution from this step is then used as an initial guess for the next step.
- Newton-Raphson step: Iterations start from the approximate solution furnished by the previous step. At any given iteration, derivative information is used to accelerate convergence towards steady-state solutions. Given the quadratic convergence rate of Newton’s method, this step can provide highly accurate solutions with much fewer iterations than the previous step. However, this procedure can also diverge if the initial guess obtained in the previous step does not lie in the convergence region of Newton’s method. Therefore, it is not robust with respect to the choice of initial guess.

### Constructing Steady-State Solution Branches

Constructing the solutions of Eq. (S19) using the computational procedure introduced in the previous section is generally time-consuming, rendering parameter-sweep computations challenging to perform. Therefore, we use Keller’s arc-length continuation method [18] to construct steady-state solution branches. The goal is to compute parametric solutions 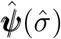 of Eq. (S23) efficiently by leveraging a predictor-corrector scheme so as to avoid the computational costs associated with repeated execution of the foregoing pseudo-linear step—the computational bottleneck of the procedure outlined in the previous section. Here, we seek 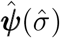 as parametric solutions of the problem

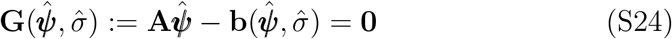

subject to

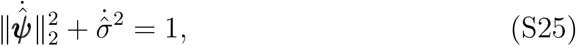

where overdot denotes differentiation with respect to the arc-length *s*. The predictor step in branch-continuation methods requires the tangent vector 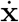, where 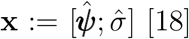 [18]. The derivatives with respect to *s* are ascertained by solving

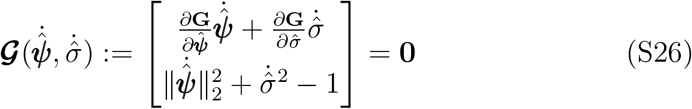

using Newton’s method. Suppose that the vector of steady-state solutions **x**_*n*_ at the *n*th arc-length step along the solution branch is known with *s*_*n*_ the corresponding arc-length. At this step, the tangent vector 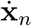 can readily be computed by solving Eq. (S26). The goal now is to compute the solution vector at the next step **x**_*n*+1_ corresponding to *s*_*n*+1_ = *s*_*n*_ + Δ*s* for a prescribed Δ*s*. First, an auxiliary function is introduced

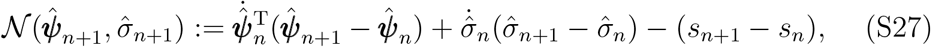

which is a linearized version of Eq. (S25). Next, **x**_*n*+1_ is computed by solving

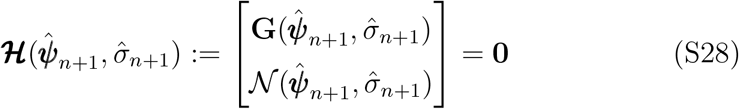

using a predictor-corrector scheme. In the predictor step, the tangent vector 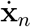 is used to construct an initial guess for **x**_*n*+1_ as follows

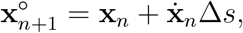

which is accurate to first order in Δ*s*. In the corrector step, **x**_*n*+1_ is computed to high accuracy by solving Eq. (S28) using Newton’s method with 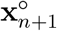 as an initial guess. A key advantage of predictor-corrector approaches is that, the initial guess generated in the predictor step usually lies in the convergence region of Newton’s method even for moderately sized Δ*s*. Moreover, constructing the initial guess in the predictor step is computationally much less costly than the pseudo-linear step discussed in the previous section. Therefore, parameter-sweep computations can be performed much more efficiently using branch-continuation methods than the procedure introduced in the previous section, if it were to be executed at all points along the solution branch.

### Stability of Steady-State Solutions

We determine the stability of steady-state solutions using linear stability analysis. Let 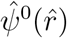 and 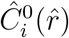 denote the steady-state solutions of Eqs. (S12) and (S13). Upon perturbations, the time-varying solutions of Eqs. (S12) and (S13) can be expressed as

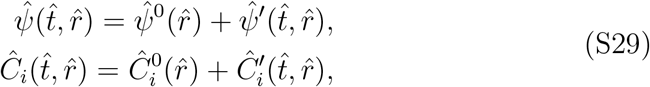

where 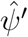 and 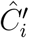 are infinitesimal perturbations induced in the protocell by fluctuations in environmental conditions. To simplify the analysis, we assume that these fluctuations can only destabilize the concentration distributions in the cell and membrane without affecting the steady state of the ocean. Accordingly, the concentration and electric-potential boundary conditions on *A*^out^ are not influenced by these perturbations. We further assume that instabilities are mainly caused by concentration perturbations, neglecting the disturbances that they can induce in the electric potential field (*i.e.*, 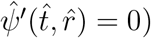). Thus, we consider the following perturbation *ansatz*

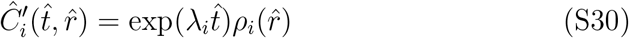

with *λ*_*i*_ the eigenvalue characterizing the dynamics of species *i*. Substituting Eqs. (S29) and (S30) in Eq. (S13), taking into account all the assumptions discussed above and neglecting the second- and higher-order terms in 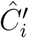, we arrive at

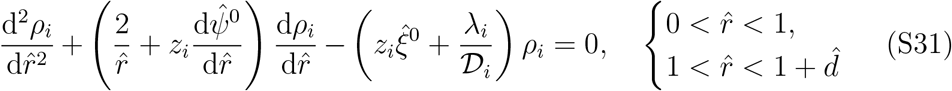

subject to

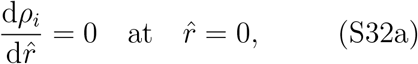

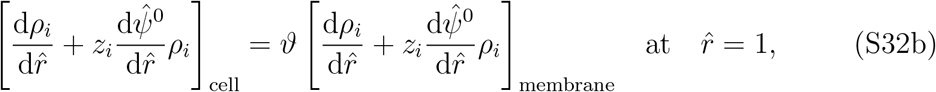

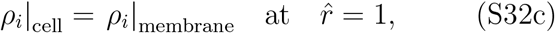

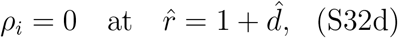

where 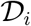 is a diffusivity coefficient defined as

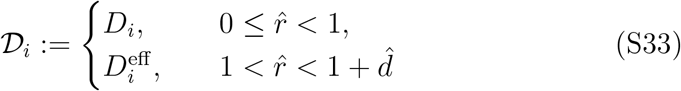

with *D*_*i*_ the the bulk diffusivity of species *i* in water and 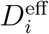 its effective diffusivity in the membrane. In our protocell model, the membrane is assumed to have a porous structure made of minerals, the effective diffusivity of which can be expressed as 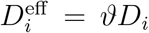 with *ϑ* the tortuosity coefficient [19, 20].

Using these definitions, the boundary condition Eq. (S32b) is derived from Eq. (S15b).

Equations (S31) can be recast into the following Sturm-Liouville form for each *i*

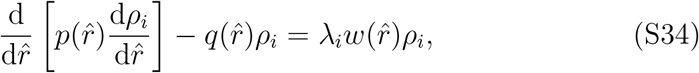

where

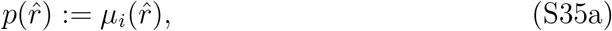

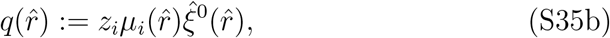

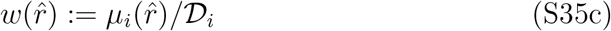

with

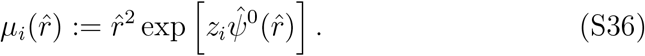

The Sturm-Liouville problem Eq. (S34) is called regular if it is subject to some variants of homogeneous Robin boundary conditions, 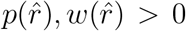, and 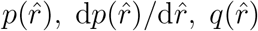, and 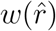 are continuous on 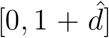 [35]. The following properties of regular Sturm-Liouville problems are of particular relevance to linear stability analysis [36]:

- Eigenvalues are discrete and real.
- Eigenvalues are bounded from above.
- Eigenfunctions form a complete orthogonal basis for an *L*_2_ Hilbert space.

Moreover, the solutions of a regular Sturm-Liouville problem are continuously differentiable [36]. However, the Sturm-Liouville problem that arise from Eq. (S31) is not regular. Firstly, the boundary condition at 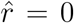 is singular because *p*(0) = 0. Secondly, 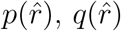, and 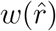 are nonsmooth at 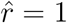. Nonetheless, modern treatments of the Sturm-Liouville theory allows these functions to satisfy more relaxed conditions, such that the foregoing three properties still hold. Accordingly, it suffices for 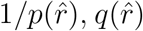, and 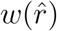 to be locally Lebesgue integrable—a condition satisfied by Eq. (S35). Although, the solutions (*i.e.*, eigenfunctions here) may satisfy weaker smoothness and continuity properties. This generalization followed from the important finding that Hilbert function spaces can be decomposed into mutually orthogonal singular and absolutely continuous subspaces for self-adjoint operators (see the work of Zettl [35] for more details).

**Figure S2:**
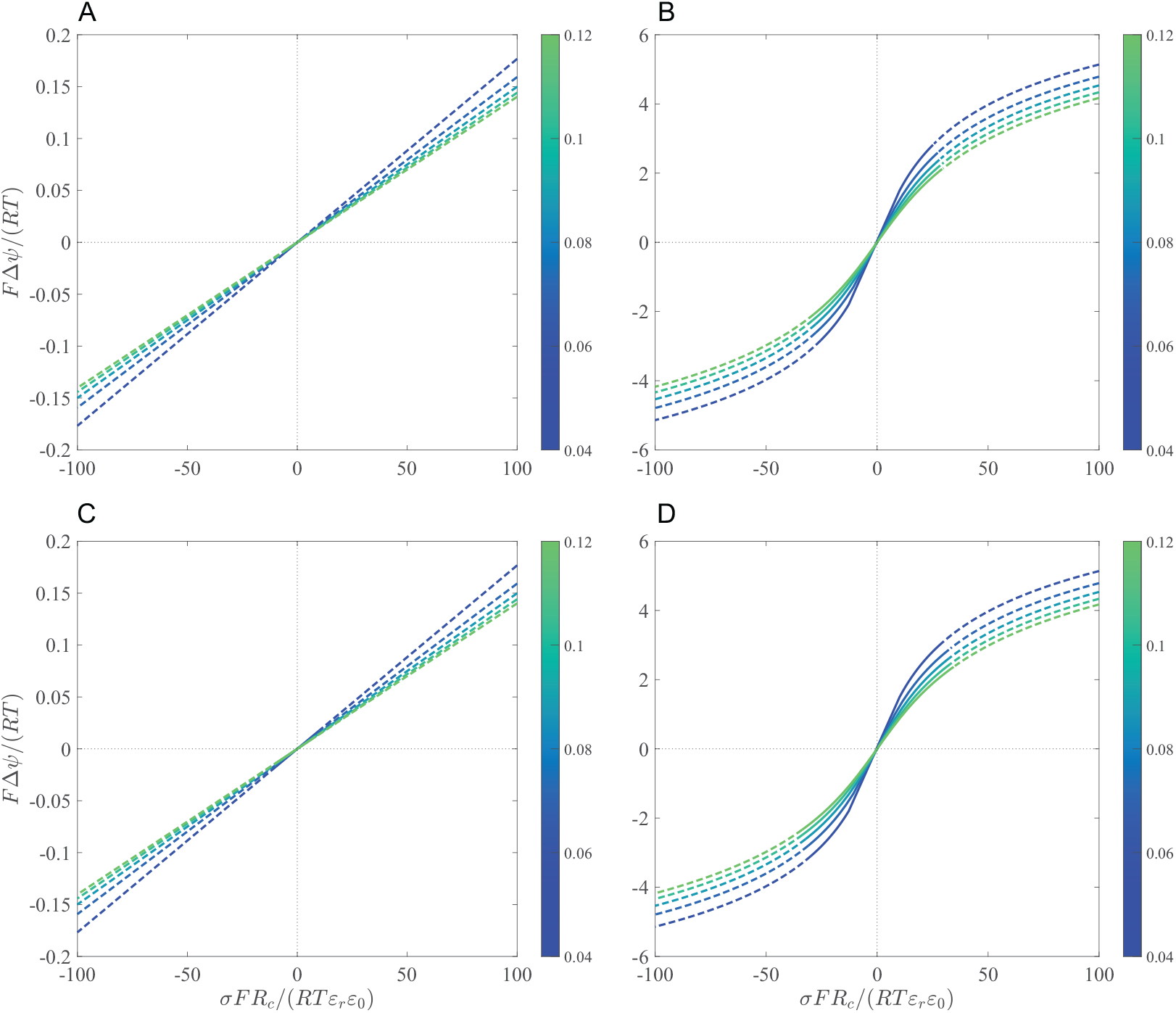
Stability along steady-state solution branches of Δ*ψ* parametrized with respect to the surface charge density *σ* at *σ*_*r*_ = 0.02 with (A) *R*_*c*_ = 10^−6^ m and *ϑ* = 0.05, (B) *R*_*c*_ = 10^−8^ m and *ϑ* = 0.05, (C) *R*_*c*_ = 10^−6^ m and *ϑ* = 0.1, and (D) *R*_*c*_ = 10^−8^ m and *ϑ* = 0.1. Colorbars indicate the value of *C*^salt^ = *C*_*∞*_/2 that corresponds to each curve in (A)–(D). The tortuosity coefficient *ϑ* only affects stability without altering steady-state solutions.

Given the properties of Sturm-Liouville problems discussed above, it suffices to show that the maximum eigenvalue of Eq. (S31) subject to the boundary conditions Eq. (S32a)–(S32d) is negative to prove that a steady-state solution is stable. We, thus, solve Eq. (S31) numerically using finite-difference methods following a similar procedure as discussed before (see “Numerical Approximation of Steady-State Solutions”). Discretization of Eq. (S31) subject to Eq. (S32a)–(S32d) results in the following systems of equations

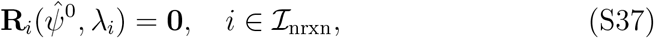

which have a nontrivial solution if and only if 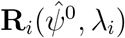 are singular. The determinant can be used as a measure of how far a matrix is from being singular, the application of which leads to the following condition

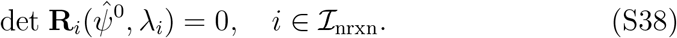

Determining matrix singularity is computationally expensive. Therefore, we consider an alternative condition for 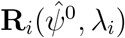 to be singular by requiring its minimum singular value to vanish

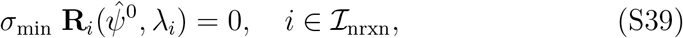

where *σ*_min_ is an operator returning the minimum singular value of **R**_*i*_ (not to be confused with the surface charge densities introduced in previous sections). Solving Eq. (S39) is computationally less expensive than solving Eq. (S38) since the singular values of **R**_*i*_ can be efficiently computed by leveraging its sparsity structure. Once Eq. (S39) has been solved, the stability of steady-state solutions can be determined by the sign of 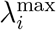 for all 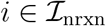, where 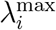 is the maximum eigenvalue of 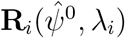 (see Figs. S2 and S3).

### Concentration Heterogeneity and Reaction Efficiency

So far, we restricted our analysis to electric potential fields that could have been indued by nonuniform distributions of inorganic ions of the primitive ocean (*i.e.*, nonreactive species). These nontrivial potential fields could have affected the operation and evolution of early metabolic cycles. Many of the organic molecules, reducing agents, and energy sources participating in these metabolic reactions would have been negatively charged, the transport of which would have been altered by the background electric potential field arising from the inorganic ions. Therefore, the resulting concentration distribution of these reactive species would have been heterogeneous, which, in turn, would have adversely affected the efficiency of early metabolic reactions. In this section, we study this phenomenon by solving diffusion-reaction mass-balance equations for reactive species that are subject to a prescribed background electric potential field in the cell.

**Figure S3:**
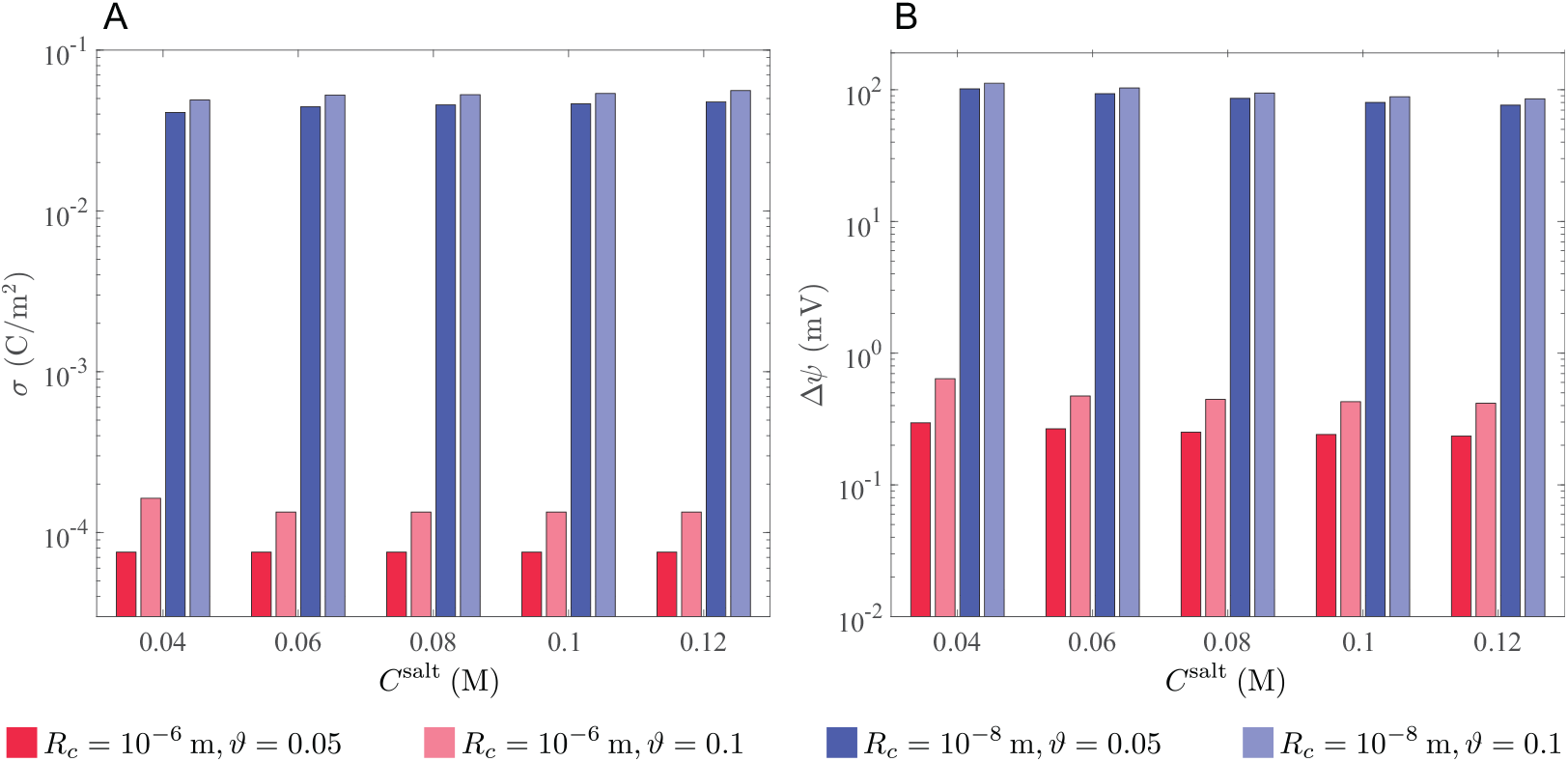
Stability limits along steady-state solution branches of Fig. S2 in the positive orthant. (A) Surface charge density, at which stability is lost. (B) Membrane potential, at which stability is lost.

Unlike nonreactive species, the steady-state fluxes of reactive species are generally nonzero. Hence, the solution strategy that we previously discussed (see “Steayd-State Solutions”) for nonreactive species is not applicable here. The concentration distribution of reactive species is also not described by Eq. (S18). Therefore, we approximate the steady-state solutions of Eqs. (S12) and (S13) for reactive species using perturbation techniques. The goal is to quantify the extent to which reaction efficiencies are affected by the background electric potential field in the cell. In the following, we first describe a quantitative measure of reaction efficiencies.

Suppose that *B* is a negatively charged reactive species and a substrate consumed by metabolic reactions taking place in the cell, which is to be imported from the ocean into the cell (*e.g.*, reducing agents or energy sources). To maximize the rate of metabolic reactions, the concentration of *B* in the cell must be maintained at the highest possible level. A positive membrane potential can enhance the transport rate of *B* from the ocean to the cell, increasing its concentration at the inner surface of the membrane 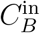, which could potentially enhance the rates of metabolic reactions. However, the overall consumption rate of *B* in the entire volume of the cell depends on its concentration distribution. A uniform distribution 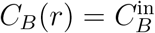 would ensure that *B* is maximally utilized by the metabolic reactions in the cell. However, uniform concentration distributions are generally not achievable due to local consumption of *B* and the background electric potential field. To quantify how concentration distributions can affect the overall consumption rate of *B*, we study a macroscopic description of its reaction-diffusion mass balance by examining the integral form of Eq. (S13). Integrating Eq. (S13) over the volume of the cell for *B* and applying the divergence theorem result in

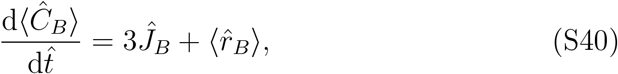

where

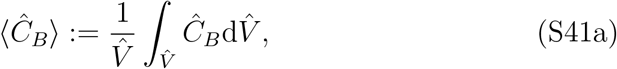

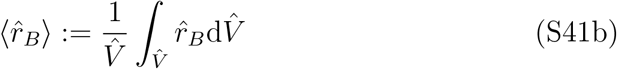

with 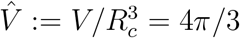. We refer to 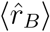 as the apparent production rate of *B*, which is a negative number here because it is consumed by metabolic reactions. The rate, at which the products of the reactions that *B* participates in are generated is proportional to −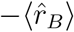. Clearly, concentration distributions that maximize 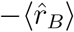 favor the progress of these metabolic reactions. To quantify the extent to which concentration distributions can enhance the overall rates of these metabolic reactions, we compare 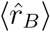 for a given concentration distribution to what it would be if *B* was uniformly distributed in the cell—the ideal distribution that maximizes its utilization. Accordingly, we define the following reaction efficiency for the consumption of *B*

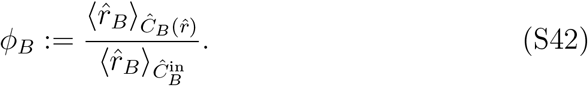

The relationship between 〈*r*_*B*_〉 and 〈*C*_*B*_〉 for nonlinear rate laws is not straight-forward. Hence, we assume that *B* is consumed in the cell according to the first-order rate law *r*_*B*_ = −*kC*_*B*_ to simplify the analysis. The apparent reaction rate from this rate law is also linear with respect to the average concentration, that is 〈*r*_*B*_〉 = −*kC*_*B*_〉. Accordingly, the reaction efficiency with respect to this rate law is

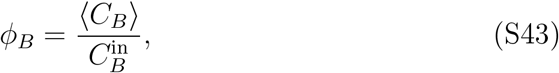

which we use as a measure of how much concentration heterogeneity can diminish or enhance the rates of metabolic reactions consuming *B*. Note that 0 *< ϕ*_*B*_ ≤ 1 only when *B* is negatively charged. However, when *B* is positively charged, *ϕ*_*B*_ can be greater than one.

Next, we compute the steady-state solutions of Eq. (S13) for *B* using finite-difference techniques along the solution branches shown in Fig. 3. These solution branches represent the steady states of the electric potential field induced by the inorganic ions of the ocean parametrized with the surface charge density. We express the dimensionless reaction rate 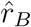 with respect to the Thiele modulus Λ_*B*_ and construct the concentration distribution of *B* in the cell that arise from the first-order rate law *r*_*B*_ = −*kC*_*B*_ by solving

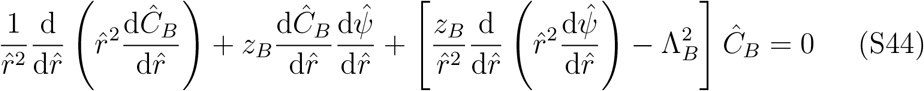

subject to

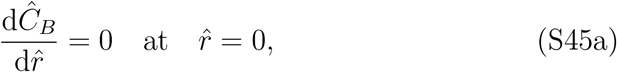

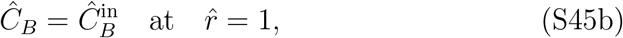

where

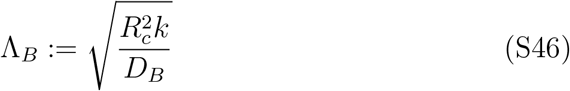

is the Thiele modulus [37].

Once the concentration distribution of *B* has been determined, we compute the reaction efficiency from Eq. (S43). As expected, the reaction efficiency for positively charged species is higher than for negatively charged ones because these ions must diffuse through a negatively charged medium to participate in metabolic reactions that occur in the cell (Fig. S4). Higher surface charge densities cause more negative ions to accumulate in the cell, amplifying this effect. The reaction efficiency is always less than one for negatively charged species. However, it can exceed one for positively charged species if the surface charge density is large enough. Note that, the background charge induced by the inorganic ions of the ocean is not the only parameter affecting the reaction efficiency. The local consumption of reactants can also result in heterogeneous concentration distributions, irrespective of the background charge. This effect is more conspicuous in the limit *σ* → 0. Even though the entire volume of the cell is electroneutral in this limit, the reaction efficiency can be less than one (Fig. S4). Note also that, diminished reaction efficiencies as a result of local mass sinks is more pronounced at larger Thiele moduli.

**Figure S4:**
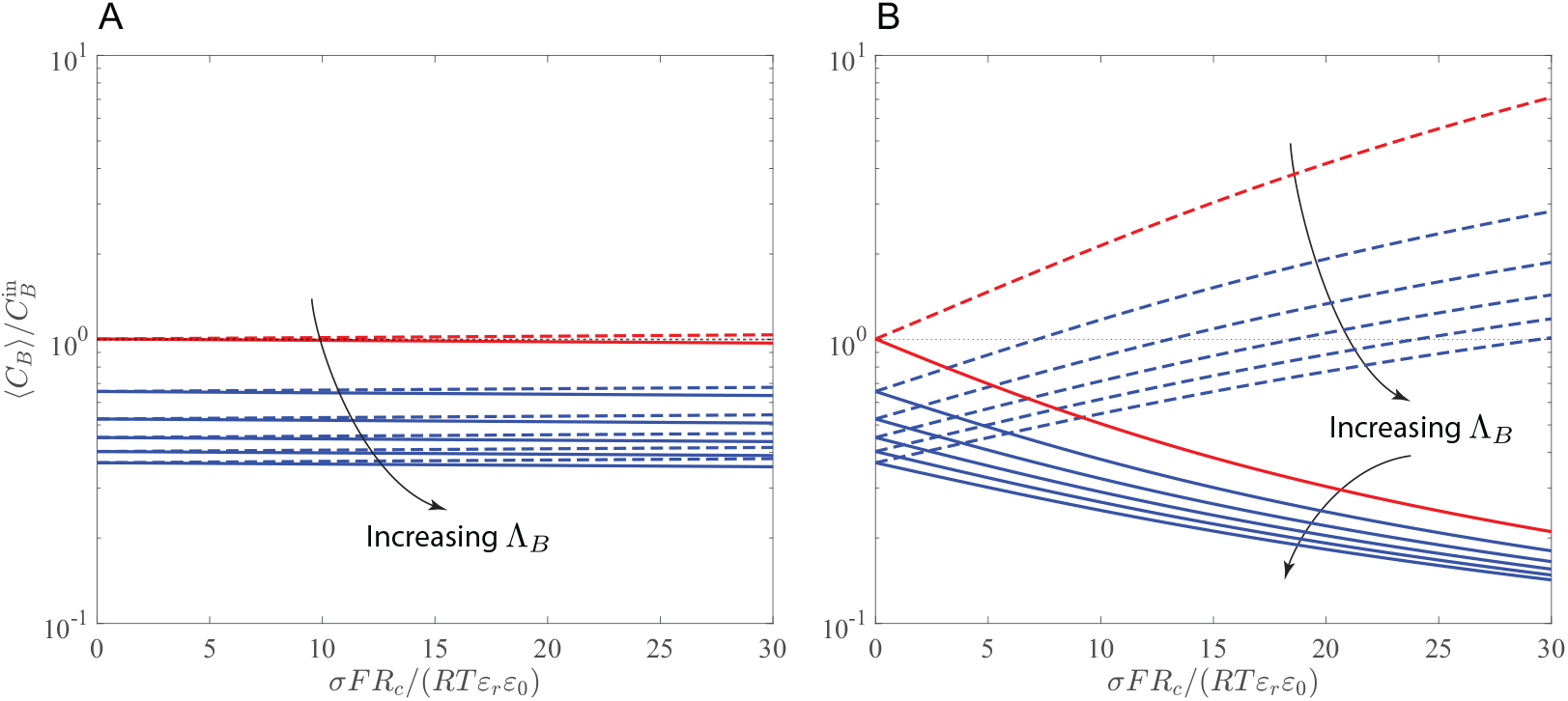
Average concentration of the cation (dashed lines) and anion (solid lines) arising from the dissociation of a monovalent salt inside the cell at *C*^salt^ = 0.1 M, 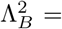 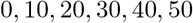, and (A) *R*_*c*_ = 10^−6^ m and (B) *R*_*c*_ = 10^−8^ m. Each Thiele modulus Λ_*B*_ corresponds to a solid-dashed curve pair, increasing along the direction indicated by the arrows. Red lines represent the nonreactive limit, where Λ_*B*_ → 0.

### Electroneutrality and Structural Stability of Protocells

Electroneutrality is often treated as a fundamental law governing the state of electrolyte systems [29]. It is also regarded as a fundamental constraint that biological systems are subject to [25, 11]. As such, it is believed to underlie regulatory responses to several stress conditions [24, 11]. However, as discussed in the main text, violation of electroneutrality is essential for the mechanism that we proposed to promote the evolution of early metabolic cycles in primitive cells that lack lipid membranes and enzymes. To corroborate this mechanism, we examine the possibility that electroneutrality was not a fundamental constraint at the earliest stages of evolution. The goal is to understand whether electroneutrality could have resulted from the evolution of lipid membranes, specialized ion channels, and active transport systems selected for to minimize catastrophic events due to osmotic crisis. From this perspective, electroneutrality is an emergent property of evolving systems, self-optimizing towards a state of maximal structural stability through natural selection.

Here, we quantitatively examine the relationship between osmitic crisis and electroneutrality through a simplified case study. We consider an electrolyte, comprising a cation *M*^+^ and an anion *X*^−^. Our objective is to determine how the osmotic pressure arising from this system varies with the charge density of the solution and assess if the minimum osmotic pressure is attained when the solution is electroneutral. There are two main variables that determine the osmotic coefficient of an electrolyte, namely the (molal) ionic strength *I*_*m*_ and the total (molal) concentration *m* of the solution [29]. The idea is to focus solely on the role of electroneutrality and identify the (molal) charge density *ξ*_*m*_ that minimizes the osmotic coefficient *φ* at fixed *I*_*m*_ and *m*.

The state of single electrolyte systems *MX* at fixed temperature and pressure is specified by two variables, namely the molal concentrations *m*_*M*_ and *m*_*X*_. Hence, specifying *ξ*_*m*_, *I*_*m*_, and *m* for a general single electrolyte system overdetermines its state. However, for a monovalent electrolyte, such as the one we considered here, *I*_*m*_ = *m/*2. Thus, the charge density can freely vary without causing an inconsistent degree of freedom. We begin by stating the virial expansion of the excess Gibbs energy of mixing

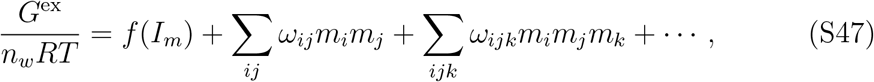

which is the basis of Pitzer’s model, where *n*_*w*_ is the mass of water in kg [29]. The first term in Eq. (S47) captures long-range electrostatic forces and the rest capture medium- and short-range interactions among ions. To simplify the analysis, we omit the terms corresponding to interactions among three or more ions as they are negligible in most cases [38]. Differentiating the excess Gibbs energy with respect to *n*_*w*_ yields the osmotic coefficient

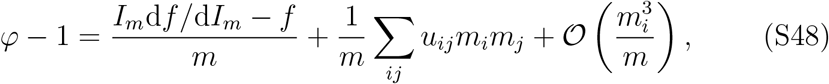

where

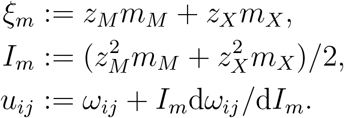

To make the analysis more concrete, we consider a case, where the electrolyte *MX* is contained in a protocell lying at the bottom of the primitive ocean, such as that shown in Fig. 1. The osmotic pressure differential across the cell membrane is derived from Eq. (S48)

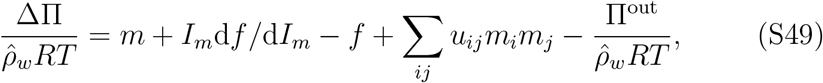

where ΔΠ ≔ Π^in^ Π^out^ is the osmotic pressure differential, Π^in^ pressure in the cell, Π^out^ pressure in the ocean, and 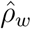 reduced water density (see the work of Akbari et al. [11] for definition and detailed discussion). We assume that the thermodynamic state of the ocean is specified, so that the last term in Eq. (S49) is a constant. In this equation, only the term corresponding to the second virial coefficient of Eq. (S47) on the right-hand side depends on *ξ*_*m*_. Therefore, it is the only variable term, with respect to which ΔΠ is minimized. Accordingly, we seek *ξ*_*m*_ that minimizes

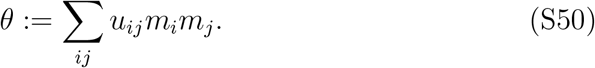

For the monovalent electrolyte *MX*, the concentration of ions can be expressed with respect to *m* and *ξ*_*m*_ as

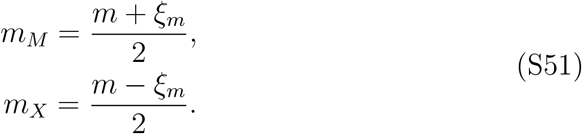

Substituting Eq. (S51) in Eq. (S50) and nondimensionalizing the resulting terms yields

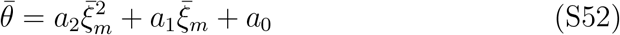

where

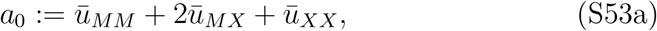

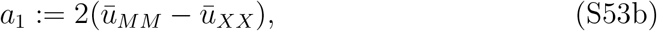

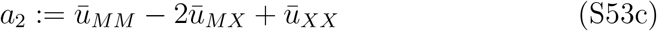

with 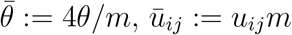, *ū*_*ij*_ ≔ *u*_*ij*_*m*, and 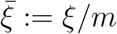.

Next, we leverage the properties of the coefficients *a*_*i*_ in Eq. (S53) that can be deduced from experimental observations. First, thermodynamic mixing properties are not significantly affected by the diagonal second virial coefficients for most electrolyte systems, so that *ω*_*MM*_ = 0 and *ω*_*XX*_ = 0 [39, Section 2.5], which, in turn, results in *ū*_*MM*_ = 0, *ū*_*XX*_ = 0, and *a*_1_ = 0. Second, *a*_0_ < 0 for a wide range of dilute electrolytes (*I*_*m*_ ≲ 0.25 mol/kg-w) [29, Eq. (50) and Table 1], from which it follows that *a*_2_ > 0. One can deduce from these empirical properties and the functional form of 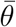 in Eq. (S52) that 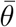 has a minimum, and it is attained at 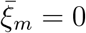.

Finally, we emphasize that the results presented in this section for a single monovalent electrolyte cannot be regarded as a rigorous proof. Nevertheless, they support the hypothesis that electroneutral systems are subject to minimal osmotic stress. More analyses are required to generalize these results to mixed electrolytes with polyvalent ions. Given the role of electrostatic forces in short- and medium-range ion-ion interactions, it may be plausible to assume that the nonlinear proportional relationship between the osmotic pressure differential ΔΠ and absolute charge |*ξ*_*m*_| is generalizable to more complex electrolyte systems. However, whether the minimum osmotic pressure is always attained exactly at 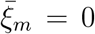, regardless of the molecular characteristics of the ions involved, warrants further investigations.

### Surface Charge of Minerals

The mechanism we introduced in this paper to generate positive membrane potentials hinges on porous membranes with positively charged surfaces. In this mechanism, the membrane potential is larger if the surface charge density on the inner surface is larger than on the outer surface of the membrane. From our case studies, we found that Δ*ψ* ~ 100 mV can be achieved in small protocells with radius *R*_*c*_ ~ 10^−8^ m for Δ*σ* ~ 0.1 C/m^2^ (see Fig. S3), where Δ*σ* ≔ *σ*^in^−*σ*^out^ is the surface-charge-density differential across the membrane. In this section, we describe a specific scenario based on experimental measurements of the surface charge density for how such positive surface charges could have been realized in primitive cells.

Solid surfaces, such as those of minerals, can adsorb or desorb ions (usually H^+^, OH^−^, or other ions that may be present in the system) from or to water when exposed to an aqueous phase. As a result, these surfaces may acquire a surface charge. When the temperature, pressure, and ionic composition of the electrolyte that mineral surfaces are subject to are specified, the surface charge density is a function of pH. At fixed temperature and pressure, the functional form of *σ*(pH) for each mineral depends on its constituents and the composition of the electrolyte, which can typically be represented by a monotonically decreasing function, such as those shown in Fig. S5 [40, Chapter 1]. However, non-monotonic *σ*(pH) have also been observed (for example, see Fig. 6 in the work of Nyamekye and Laskowski [41]). Nevertheless, we only focus on minerals that exhibit a monotonic *σ*(pH) in this section.

**Figure S5:**
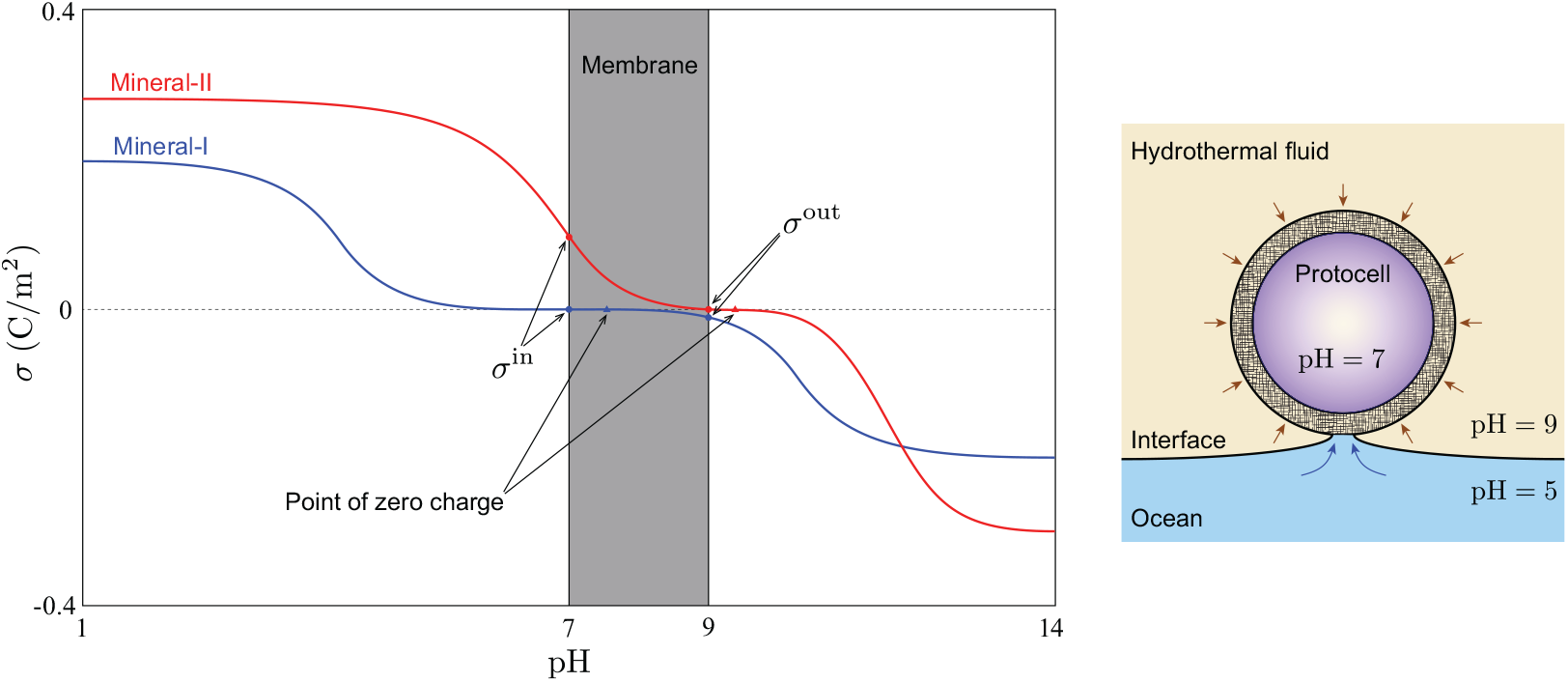
Typical surface charge density of minerals measured as a function of pH using potentiometric-conductometric titration experiments [40, Chapter 1]. A pH gradient across the membrane of the protocell model shown in Fig. 1 is assumed to cause a surface-charge-density differential between the inner and outer surfaces of the membrane. The protocell resides near a hydrothermal vent at an interface between two fluids with pH ≈ 9 and pH ≈ 5. The fluids flow into the cell from the alkaline vent and acidic ocean. The acidic and alkaline fluids neutralize into water, such that pH ≈ 7 in the cell. Mineral-I and Mineral-II are two hypothetical minerals that exhibit different functional forms for *σ*(pH). For the given pH gradient between the hydrothermal vent and ocean, the surface charge densities formed on the inner and outer surfaces of a membrane made of Mineral-I are almost identical. However, a large surface-charge-density differential can be generated across a membrane made of Mineral-II, such that *σ*^in^ and *σ*^out^ are both positive, similarly to the scenario described in the left diagram of Fig. 1B.

The monotonicity of *σ*(pH) implies that |*σ*| attains its maximum in the alkaline and acidic limits. Accordingly, surface charge densities for most minerals are observed in the range *σ*(pH = 1) *< σ < σ*(pH = 14). For example, surfaces charge densities in the range −0.4 *< σ <* 0.4 C/m^2^ for synthetic and natural ferrous minerals, which are relevant to the conditions on the primitive Earth [5, 42], have been reported [43, 44] with a similar *σ*(pH) to those shown in Fig. S5. This range generally agrees in order of magnitude with the stable ranges of *σ* that we ascertained in our case studies (see Figs. S2 and S3).

Alkaline hydrothermal vents are considered to be one of the likely environments, in which life could have originated [8, 1]. In these environments, CO_2_-rich acidic ocean water (pH ≈ 5) could have interfaced with alkaline hydrothermal fluids (pH ≈ 9), providing suitable conditions for the first metabolic reactions to emerge [42]. Recently, experimental evidence has been found, suggesting that such pH gradients could have provided sufficient energy to drive the thermodynamically unfavorable carbon-fixation steps of early metabolism under prebiotic conditions [45]. Here, we suggest that the pH gradient between hydrothermal fluids and the primitive ocean could also have generated sufficiently large surface-charge-density differentials across protocell membranes (Δ*σ* ~ 0.1 C/m^2^).

To clarify the point raised above, consider a protocell, residing at an interface between a hydrothermal vent and ocean (Figs. S5). Suppose that acidic and alkaline fluids flow into the cell, neutralizing each other, such that pH ≈ 7 in the cell. The neutral pH in primitive cells would have been optimal for the emergence of surface metabolism at the origin of life [5]. The difference between *σ*^in^ at pH = 7 and *σ*^out^ at pH = 9 may be large or small, depending on the constituent minerals of the membrane. Mineral-I and Mineral-II in Fig. S5 represent two hypothetical minerals that could generate small and large Δ*σ*, respectively. A key difference between the functional form of *σ*(pH) for these minerals is in the point of zero charge (PZC) [40, Chapter 1]. The PCZ occurs at pH ≈ 7.5 for Mineral-I. Thus, *σ*^in^ and *σ*^out^ are both small, so that Δ*σ* ≈ 0 C/m^2^. However, for Mineral-II, the PCZ occurs at pH ≈ 9.3. As a result, *σ*^in^ and *σ*^out^ are both positive with Δ*σ* ≈ 0.1 C/m^2^. Therefore, the scenario we described in Fig. 1B for generating positive membrane potentials could have been realized for protocell membranes made of Mineral-II. Interestingly, experimental measurements of the surface charge density of mineral-water interfaces indicate that the functional from of *σ*(pH) for several transition-metal sulfides (*e.g.*, Ni_3_S_2_ and ZnS) is similar to that of Mineral-II in Fig. S5 with the PZC in the alkaline range [41, 46].

## Acknowledgments

This work was funded by the Novo Nordisk Foundation (Grant Number NNF10CC1016517) and the National Institutes of Health (Grant Number GM057089).

## Author Contributions

Conceptualization, A.A. and B.O.P.; Methodology, A.A.; Validation A.A.; Formal Analysis A.A.; Investigation, A.A.; Writing – Original Draft, A.A. and B.O.P.; Writing – Review & Editing, A.A. and B.O.P.; Funding Acquisition, B.O.P.; Resources, B.O.P.; Supervision, B.O.P.

